# Facultative winter accession of *Camelina sativa* (L. Crantz) with early maturity contributes to understanding of the role of *FLOWERING LOCUS C* in camelina flowering

**DOI:** 10.1101/2022.05.30.494064

**Authors:** Matthew A. Ott, Ratan Chopra, Katherine Frels, Anthony Brusa, Eva Serena Gjesvold, M. David Marks, James A. Anderson

## Abstract

Camelina is being developed as a winter oilseed cover crop. Early flowering and maturity are desired traits in camelina to allow for relay planting or seeding of a summer annual following camelina harvest. Here we report that while all winter biotype accessions of camelina have a functional allele of *FLOWERING LOCUS C (FLC)* on chromosome 20, there are also at least 20 previously characterized spring biotype accessions that have a functional *FLC* allele at this locus. We observed this by analyzing 75 accessions (67 spring type, one facultative, and seven winter type) that were resequenced by Li et al., (2020) as well as 21 additional accessions for this analysis. This discovery will inform marker assisted selection efforts that are underway to increase genetic variation in the genetically narrow base of winter camelina germplasm. Furthermore, we optimized a KASP genotyping approach that effectively differentiates the presence of either the functional or subfunctional *FLC* allele on chromosome 20. These analyses identified a facultative winter biotype accession of camelina (PI650163-1, winter hardy with subfunctional chromosome 20 *FLC* allele) that has demonstrated two years of winter-hardiness and has flowered at least a week earlier than the common winter accession, ‘Joelle’. A bioinformatics approach to cytotype analysis in camelina also provided more precise categorizing of camelina accessions in the USDA-NPGS germplasm into 2n=38 and 2n=40 cytotypes. Early maturing winter-hardy camelina will reduce stress on a subsequent soybean crop and improve total cropping system yields when camelina and soybean are grown sequentially in the same season on the same land.

## 1. Introduction

There is unprecedented global demand for biofuels and feed byproducts as consumers seek out fossil fuel alternatives to mitigate climate change and energy price volatility. In the Upper Midwest winter camelina is being developed as a new cover/cash crop that can be planted in the fall and harvested in spring on the same lands used for traditional summer crop such as that undergoing the maize to soybean rotation on much the Midwestern farm acreage. When planted as a winter annual oilseed cover crop, *Camelina sativa* (L. Crantz) should be able to ease some of the supply shortage issues for both the energy and feed sectors. Fall pllanted/spring harvested camelina as an oilseed cover crop also provides important ecosystem services, such as reducing nitrate leaching and providing supplemental nutrition for pollinators, further promoting the crop as a potential solution to several complex challenges (Weyers et al., 2019; Eberle et al., 2015).

*Camelina sativa* has been shown to be descended from *Camelina macrocarpa*, with a region of origin spanning the Mediterranean and Eastern Europe (Brock et al., 2018; Chaudhary et al., 2020). It remains unclear exactly when *Camelina sativa* began to be actively cultivated rather than being tolerated as a weed in cultivated flax (Brock et al., 2018), but its use as a crop can be documented as far back as the early Iron Age (∼1200 BCE) (Zohary et al., 2012). Two cytotypes exist for allohexaploid *Camelina sativa* – either n=6+7+7 (20) or n=6+6+7 (19) (Chaudhary et al., 2020; Hotton et al., 2020; Brock et al., 2022). The genome size was estimated to be 785 Mb, which is relatively small for a hexaploid species (Kagale et al., 2014). Camelina is closely related to Arabidopsis thaliana as both are in Brassicacea lineage group I (Huang et al., 2016). Similar to Arabidopsis, camelina is largely a self-fertilized species. In the *Camelina sativa* germplasm within the US’s National Genetic Resources Program, only eight of the 41 accessions exhibited the winter biotype, while the rest exhibited the spring biotype and are not winter-hardy (Hotton et al., 2020). This leaves plant breeders focused on winter types for their value as a winter cover crop with a narrow genetic base. To increase genetic diversity for key traits in winter camelina, one strategy is to cross-pollinate winter and spring types, and either select for winter types in the field if feasible, or backcross as necessary to recover the winter growth habit while introducing impactful alleles. Identifying genetic markers that can predict the winter/spring growth habits in *Camelina sativa* would save time otherwise spent waiting for populations to phenotypically segregate for this trait before selection and desirable allele stacking could take place in a breeding program. In Arabidopsis, a functional *Flowering Locus C* (*FLC*) gene was found to impart a requirement for vernalization ahead of flowering, which the hallmark of a winter type. Similarly in camelina, the presence of functional or subfunctional *FLC* orthologs have been shown to effectively differentiate between two spring accessions of camelina (DH55 and CO46) and one winter accession (Ames33292, ‘Joelle’) using qRT-PCR (Anderson et al., 2018). These analyses identified a SNP in FLC on chromosome 20 that distinguishes between the spring and winter habits. We have used this information to optimize a genotyping approach that differentiates winter from spring camelina accessions based upon the chromosome 20 FLC SNP using DNA-based Kompetitive Allele Specific PCR (KASP). To expand the our information on the importance of this

SNP in controlling flowering time we resequenced 21 camelina accessions within our collection and downloaded an additional 75 resequenced accessions from data on NCBI provided by Li et al., (2020). While this *FLC* SNP helped differentiate between most spring and winter lines, we identified several spring lines with the winter type *FLC* allele. This information is being used to facilitate the breeding of winter hardy lines that flower and mature early.

## 2. Materials and Methods

### 2.1 Field and indoor screening of *Camelina sativa* germplasm

In 2015 we requested 420 accessions of *Camelina sativa* from the USDA- National Plant Germplasm System, Plant Genetic Resources of Canada, the Plant Breeding and Acclimatization Institute (IHAR) based in Poland, and Dr. Johann Vollmann at the Universität für Bodenkultur Wien (BOKU), Vienna, Austria for agronomic characterization and winter hardiness evaluation. Each accession was planted in a single, 1.5m row on September 1^st^, 2015 using a Wintersteiger Seedmech TRM 2200 plotmatic four-row cone planter with a 38 cm row spacing. Approximately 100 seeds were planted per row. On November 25^th^, 2015, each row was scored for the presence of reproductive stems or “bolting”. Accessions were scored as bolting or slow bolting.

Accessions that were scored as slow bolting were planted in triplicate in a randomized complete block design without vernalization in the greenhouse in St. Paul, MN on February 19^th^, 2016. The greenhouse temperature was maintained near 20 °C and was supplemented with halogen lighting to ensure 16h light was available each day. The height of each plant was measured 34 days after planting.

On September 13^th^, 2017 we planted a single four-row plot of each *Camelina sativa* accession previously identified as slow bolting winter types in addition to a single four-row plot USDA-NPGS accession PI650163 (origin – Former Soviet Union) in the field in Rosemount, in addition to multiple plots of the check, Ames 33292 (‘Joelle’) accession. This experiment was planted as a seed increase. The seeding rate was 0.4 g per plot and a SRES 4-row plot planter was used with a 38 cm row spacing. The seeding depth was set to 1.27 cm, however, the actual seeding depth varied and was generally deeper due to uneven soil conditions.

On October 18^th^, 2018 a selected camelina line from accession PI 650163, referred to as PI 650163-1 and the winter camelina check accession, Ames 33292 (‘Joelle’), were each planted in a 1.5m row in the field in St. Paul. PI 650163-1 was identified in the aforementioned Rosemount, MN field in May of 2018 as the earliest flowering plant present. The seeding rate was 0.1 g per row, the seeding depth was 1.27 cm and a SRES 4-row plot planter was used with a 38 cm row spacing.

On September 9^th^, 2019 camelina accessions PI 650163-1 and Ames 33292 were planted in duplicate in a St. Paul field in a completely randomized design to quantify differences in flowering time between these accessions. This site-year was planted by hand at a seeding rate of 0.1 g per 1.5m row and a target seeding depth of 1.27 cm.

### 2.2 Resequenced Camelina accessions

Twenty one camelina accessions were resquenced at the University of Minnesota (UMN) along with the 75 camelina accessions that were resequenced by Li et al., (2020) at Montana State University (MSU). The 21 UMN resquenced accessions were sequenced in four separate runs by the University of Minnesota Genomics Center with an Illumina NovaSeq 6000 platform on S1, S1, S4, S4 flow cells and yielded approximately 38x, 90x, 80x, 20x coverage, respectively.

### 2.3 Variant Calling Pipeline

For the Li et al., (2020) resequenced accessions, interleaved fastq.gz sequence files were downloaded from NCBI using Google Cloud Platform. Each interleaved fastq file was de-interleaved into two separate fastq files using a script provided by Daniel Standage on a Biostars forum post (https://www.biostars.org/p/141256/). Each of these files contained one of the two paired end read sequence results. These de-interleaved files were used for aligning to the *Camelina sativa* reference genome (Kagale et al., 2014) downloaded from the Crucifer Genome Initiative (developed by researchers in Dr. Isobel Parkin’s lab as a collaboration between Agriculture and Agri-Food Canada and the Global Food Security Institute, http://cruciferseq.ca/). UMN data was released from the University of Minnesota Genomics Center as zipped, de-interleaved files. All sequence data was aligned using the Burrows-Wheeler transformation aligner (version 0.7.17-r1188, Li and Durbin, 2009)(bwa- mem), and Samtools (version 1.14, Danecek et al., 2021) was used for sorting and compressing the elements of each file. Variants were called using BCFtools (version 1.9, Danecek et al., 2021) and the utilities mpileup (using options -Ou -f) and call (using -Ou - mv).

### 2.4 Alignment of WGS reads for cytotype determination

WGS reads from 14 camelina accessions were aligned to the camelina sativa reference genome specifically for determining their cytotype as has been done previously (Chaudhary et al., 2020). This alignment protocol used Bowtie2 version 2.3.4.1 with the sensitive local parameter and the minimum alignment score parameter of 20 + 10 * ln(read length), or specifically in the Bowtie2 language: --score-min G, 20, 10. Then, Bedtools version 2.29.2 was used to count the number of aligned reads in 100 kb windows that had non-zero coverage. This coverage data was then used to generate a Circos plot using TBtools-II version 1.120.

### 2.5 KASP genotyping to distinguish functional from subfunctional alleles of FLC on chromosome 20

DNA was isolated from a small leaf piece (∼1mm x 2mm leaf sample) using Sigma- Aldrich® Extract-N-AMP DNA extraction (SKU: E7526) and dilution (SKU: D5688) solutions from MilliporeSigma Inc. (St. Louis, MO 63178). KASP primers were designed as a collaboration between us and Dr. Nisha Jain (3CR Bioscience, Enfield, England) and ordered from Integrated DNA Technologies, Inc (Table 1). Tm and binding site affinity were estimated through SnapGene Viewer. The primer mix included 3.5 µL of the HEX/Y primer (fluorescence 533-580), 4.5 µL of the FAM/X primer (fluorescence 465-510), 13 µL of the common primer, and 20 µL of nuclease free water. The PCR master mix used was 3CR Bioscience’s PACE2.0 Genotyping Master Mix 2x low ROX (003-0006), and was combined with the primer mix at the following rate per reaction: 0.15 µL primer mix, 5 µL PACE2.0 Master Mix, 5 µL nuclease free water and 1 ul of DNA. KASP genotyping PCR settings on a Roche LightCycler® 480 were:

1. 95 °C enzyme activation for 15 minutes
2. A touchdown temperature cycle starting with an annealing temperature of 59°C for one minute, then a melting temperature of 95°C for 10 seconds, with a step down of the annealing temperature of 2.2°C until an annealing temperature of 51°C was reached.
3. An annealing temperature of 50 °C was used for the remainder of the PCR – all other temperatures and durations remained the same.

**Table 1.**
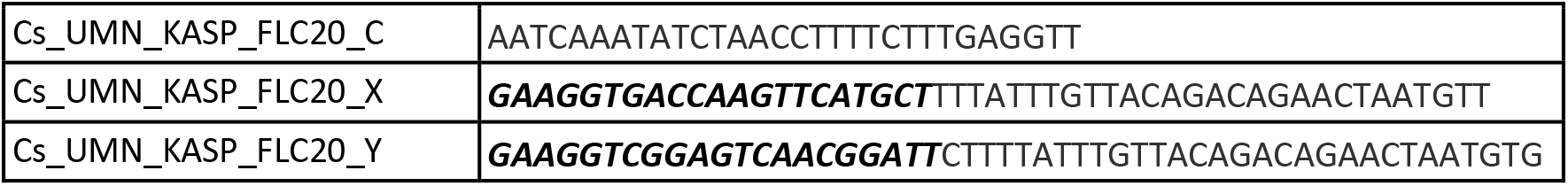
Primer design for Kompetitive Allele Specific PCR (KASP) genotyping for winter (allele X) versus spring (allele Y) growth habit in *Camelina sativa* based upon the the previously identified single basepair INDEL in exon 5 of the *FLOWERING LOCUS C* gene on chromosome 20 (Anderson et al., 2018). The bold, italic DNA sequence at the beginning of the X and Y KASP primers represent the sequences that bind to the FAM and HEX fluorescent dyes, respectively.

### 2.6 Bulked segregant analysis for flowering time

From an F_2_ population of 75 plants segregating for flowering time from a cross of PI 650163-1 and Joelle, seven early flowering plants and eight late flowering plants were selected to form DNA pools for bulked segregant analysis. DNA was first isolated from each plant individually using a Qiagen DNeasy® Plant Mini Kit, then an equal amount of DNA from each early flowering plant (180 ng) was pooled and likewise for each late flowering plant (250 ng). This DNA was submitted to the University of Minnesota Genomics Center for whole genome sequencing using NovaSeq on a S4 flow cell to achieve a target coverage of 80x.

The whole genome sequence zipped fastq files were first aligned to the *Camelina sativa* reference genome using the Burrows-Wheeler transformation aligner (version 0.7.17-r1188, Li and Durbin, 2009)(bwa-mem), and Samtools (version 1.14, Danecek et al. 2021) was used for sorting and compressing the elements of each file. Variants were called using BCFtools (version 1.9, Danecek et al., 2021) and the utilities mpileup (using options - Ou -f) and call (using -Ou -mv). The resulting VCF file was re-formatted in R using the package ‘splitstackshape’ version 1.4.8 by splitting the “FORMAT” column for both the late and early flowering pools on the characters of comma and colon to create additional columns that isolated allelic depths and genotype quality scores for each pool conforming to the format required by the R package QTLseqr (Mansfeld and Grumet, 2018, based on the G prime statistic developed by Magwene et al., 2011).

Variants were filtered using the QTLseqr package and the following parameters: refAlleleFreq = 0.20, minTotalDepth = 100, maxTotalDepth = 400, minSampleDepth = 40, minGQ = 99. The Gprime analysis in QTLseqr was then used with the following parameters: windowSize = 1e6, outlierFilter = “deltaSNP”, filterThreshold = 0.1, where the filter threshold represents a tolerance value for filtering out relatively low confidence QTL.

Finally, the plotQTLstats function was used with the following parameters: var = “negLog10Pval”, plotThreshold = TRUE, q = 0.01, where q represents the false discovery rate (FDR).

## 3. Results and Discussion

### 3.1 Discovery of a group of spring camelina accessions that are homozygous for functional *FLC* alleles on chromosome 20

In Arabidopsis the *FLC* gene is required for the winter growth habit. The *FLC* gene in Arabidopsis codes for a protein that has been shown to suppress the flowering promoter gene, *FLOWERING LOCUS T* (*FT*) (Samach et al., 2000), and is itself suppressed (methylated) by proteins that are coded by genes activated by exposure to cool temperatures (i.e. 5°C ± 1°C) (Sheldon et al., 2000). Polymorphisms in orthologous *FLC* alleles can differentiate the spring versus winter growth habit. In the winter camelina accession Joelle (Ames 33292), *FLC* is expressed on chromosomes 8 and 20, but not on chromosome 13, while its only expressed on chromosome 8 in the spring camelina accession ‘CO46’ (Anderson et al., 2018). The subfunctional allele of this gene in spring type camelina has a frameshift mutation in exon 5 on chromosome 20 at bp position 4,195,043 (Anderson et al., 2018). As we have looked at these alleles more broadly among a greater number of accessions within the camelina germplasm phenotyped for growth habit and resequenced by Li et al., (2020) (with nearly identical phenotypes reported here in Supplementary Table 1), we have found that there are at least 20 exceptional cases in which a spring accession is homozygous for the winter *FLC* allele on chromosome 20, two cases in which a spring accession is homozygous and three cases in which a spring accession is heterozygous for the winter *FLC* allele on chromosome 8, one case in which a winter accession (PI650143) is homozygous for the spring *FLC* allele on chromosome 8, and one case in which a winter accession

(CN113692) is homozygous for the spring allele on chromosome 20 (Table 2). Winter accession CN113692 was previously characterized as having a strong winter allele of *FLC* on chromosome 8 and a spring allele on chromosome 20 (Chaudhary et al., 2023). Ten additional camelina accessions in Table 2 were previously characterized for gene expression by qRT-PCR of *FLC* on chromosomes 8 and 20 and are noted as such therein (Chao et al., 2019). We found that two of these ten accessions had the spring growth habit with a winter allele of *FLC* on chromosome 20, and Chao et al., (2019) found that transcript abundance at this locus was relatively low for them, which explains the spring growth habit of these accessions and provides a clue about the underlying genetic basis of this phenomenon. Thus, when genotyping for growth habit when these exceptional spring type accessions with a winter *FLC* 20 allele are parents of a given population, the qRT-PCR method developed by Chao et al., (2019) is recommended. For the cases that involved a heterozygous genotype where homozygosity is expected in this self-pollinating species, possible explanations are that either there was a rare case of outcrossing or resequencing sample contamination.

**Table 2.**
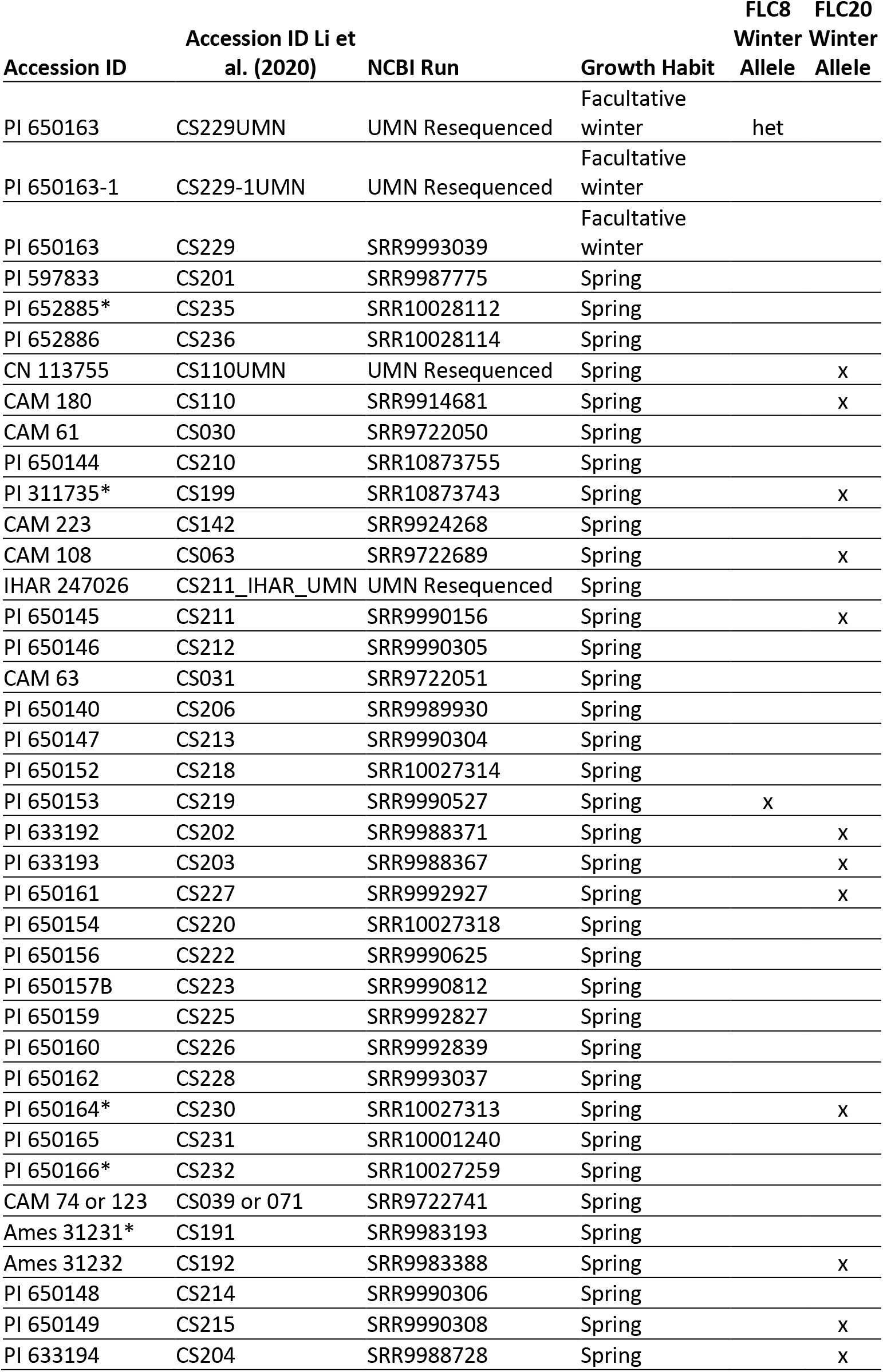
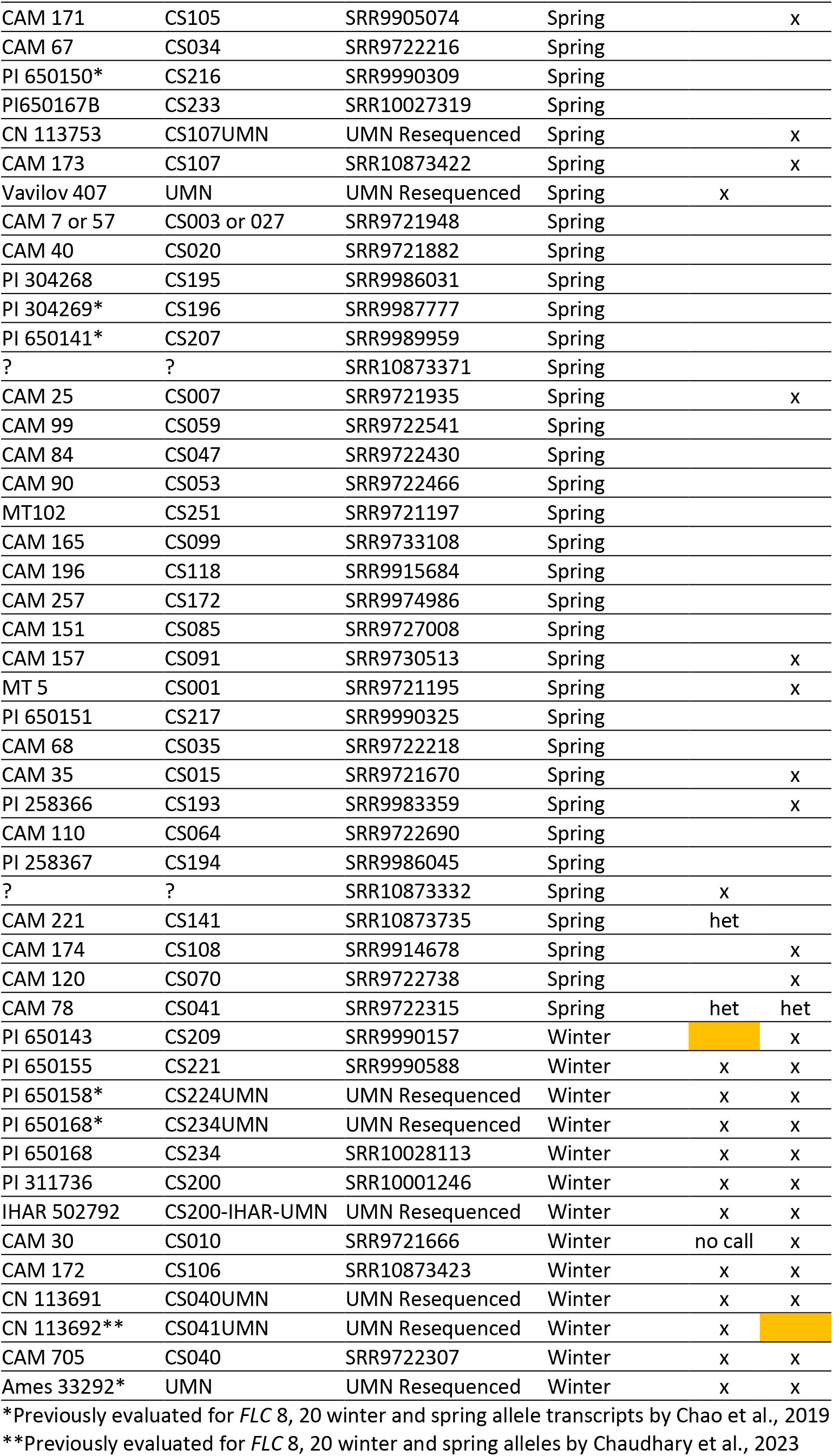
Camelina sativa accessions that were resquenced and analyzed for FLOWERING LOCUS C alleles that could predict growth habit. Resequenced by either the University of Minnesota or by Montana State University. Spring growth habit plants with winter alleles or winter growth habit plants with spring alleles are highlighted in orange.

### 3.2 Bulked segregant analysis for flowering time

The objective of this experiment was to identify the primary chromosome specific gene(s) controlling flowering time differences between the facultative winter accession of camelina PI 650163-1 and a winter accession Ames 33292 (Joelle). The QTLseqr R-package analysis identified just one major peak of QTL LOD score values (-log_10_ of variant p-values) in the bulked segregant analysis experiment (Figure 1). The variant (SNP) with the maximum LOD score (5.6) was at position 4.7 Mb on chromosome 20, was in an intergenic region and thus was unlikely to influence the flowering time trait. However, it was likely in linkage with the frameshift inducing INDEL in *FLC* on chromosome 20 (Anderson et al., 2018), which had a LOD score of 5.4 and is at position 4.2 MB. This result supports our claim that *FLC* on chromosome 20 is the primary regulator of flowering time, rather than *FLC* on chromosome 8 (for which PI 650163-1 has a subfunctional allele) or any other regulatory genes related to flowering time for PI 650163-1. There may be a few smaller signals of SNP effects on chromosomes 4, 10, and 19, but none were comparable to the signal on chromosome 20. Though the fully functional allele of *FLC* on chromosome 8 is very conserved and highly expressed among winter accessions and the subfunctional allele is conserved among spring accessions (Anderson et al., 2018), it appears that it plays a smaller role in controlling flowering time on the growth habit level in camelina, despite the fact that another report showed it has a significant effect on flowering in spring camelina backgrounds (Li et al., 2020).

**Figure 1.**
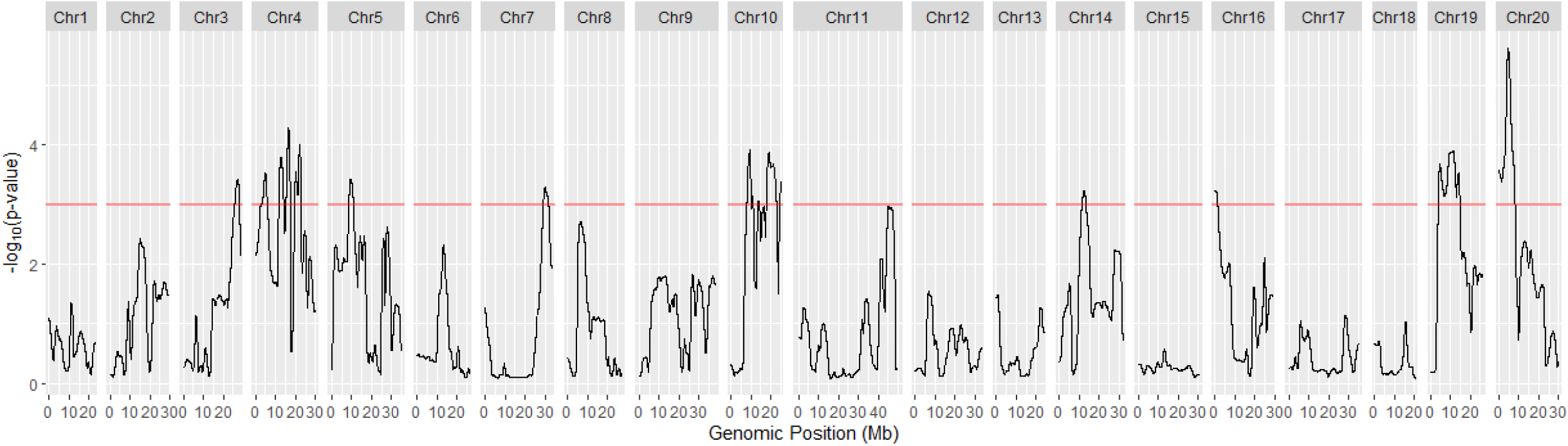
QTL mapping results obtained with QTLseqr and developed using bulked segregant analysis on an F_2_ population of a cross between Ames 33292 (Joelle) and PI650163-1. One region on chromosome 20 in *Camelina sativa* L. Crantz is a primary regulator of flowering time. The *FLOWERING LOCUS C* gene is within this region.

### 3.3 Cytotype of 14 camelina accessions in the USDA-National Plant Germplasm System

Previously ten accessions of *Camelina sativa* within the USDA-NPGS had been cytotyped as 2n=38 and one accessions as 2n=26 by a chromosome squash and counting approach (Hotton et al., 2020). This approach elucidated approximate numbers of chromosomes for these *Camelina spp.* accessions, but could not positively identify the exact *Camelina* species for them. We had made crosses among some of these stated 2n=38 and 2n=40 accessions and found that the progeny were fertile in subsequent generations (data not shown). For this reason, we included eight of these accessions in a lane for whole genome sequenced and then cytotyped bioinformatically compared to control accessions consisting of 2n=14, 2n=26, and 2n=40 as has been described previously (Chaudhardy et al., 2020). Seven of these eight we found to have the 2n=40 cytotype by the bioinformatics approach, which explained why the crosses we had made among some of these accessions were fertile (Figure 2, Supplementary Table 2). PI650167 was cytotyped as 2n=38 by Hotton et al., and we confirmed this cytotype bioinformatically. With supporting data from Chaudhary et al., 2020 and Brock et al., 2022, we were able to identify this accession as *Camelina microcarpa* along with accession IHAR50003 from the IHAR germslasm system in Poland, which lists it as *Camelina sativa*. Also with this supporting data, we were able to identify PI650152 as *Camelina rumelica* (2n=26) along with accession CN113657, which is listed as *Camelina sativa* in Plant Gene Resources of Canada.

**Figure 2.**
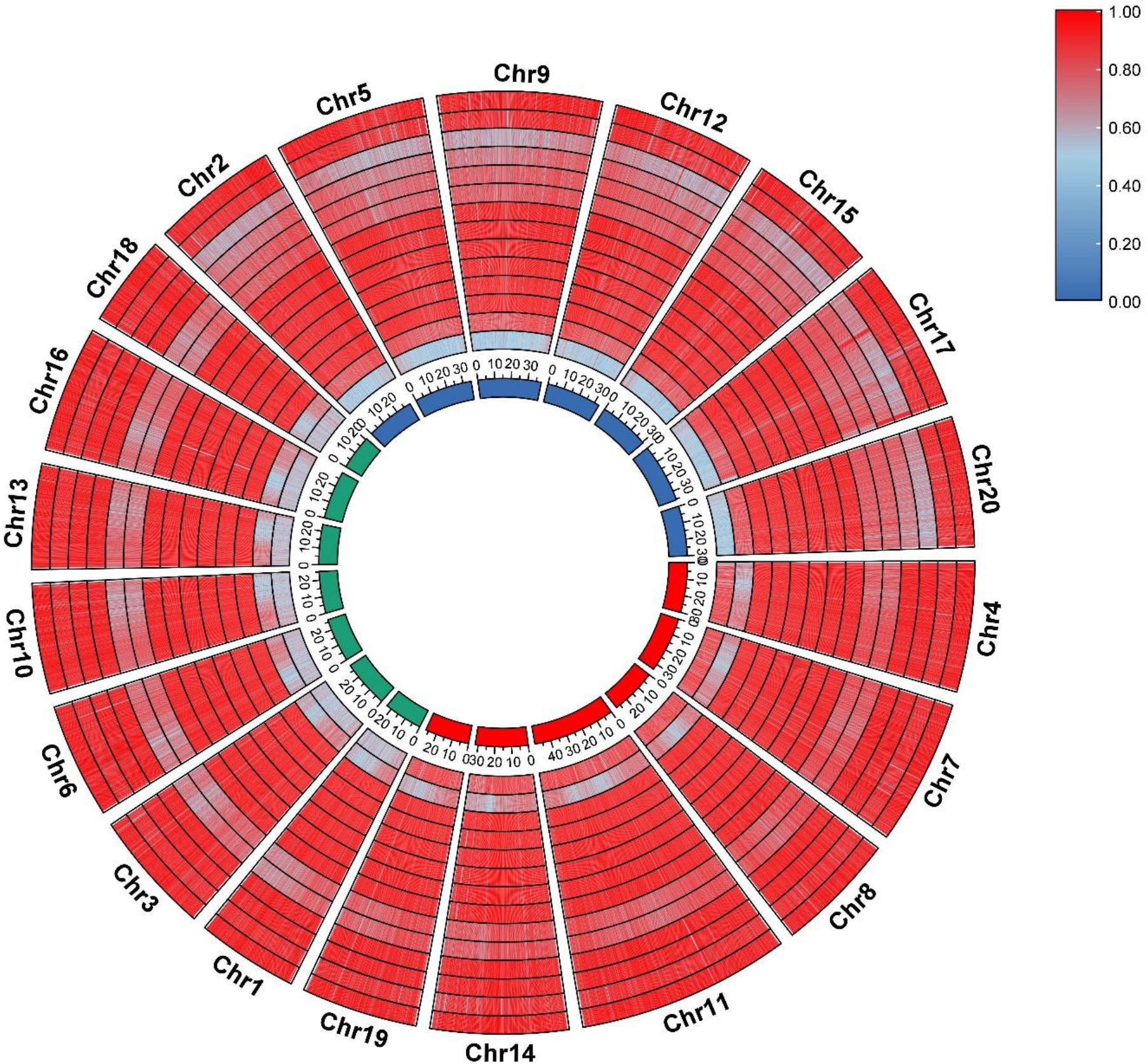
Circos plot heat map of the percentage of bases with non-zero coverage in 100kb windows from whole genome sequenced paired end reads aligned to the *Camelina sativa* reference genome. From the center to outside of the Circos plot: the three subgenomes of *Camelina sativa* – subgenome one in red, two in green, and three in blue. *Camelina spp*. accessions PI650135* (diploid *Camelina neglecta*), PI650133* (diploid *Camelina hispida*), and USDA-NPGS stated *Camelina sativa* accessions PI258367**, PI650141**, PI650143**, PI650144**, PI650146**, PI650164**, PI650152*** (*Camelina rumelica* 2n=26), CN113657 (*Camelina rumelica* 2n=26), PI650167**** (*Camelina microcarpa* 2n=38), IHAR50003 (*Camelina microcarpa* 2n=38), PI650168**, PI650163-1. *Previously cytotyped as diploid *C. spp.* by DNA sequencing by Chaudhary et al., 2020. **Previously cytotyped as hexaploid *Camelina* (2n=38) by chromosome squash by Hotton et al., 2020, but found to be hexaploid *Camelina sativa* (2n=40) by DNA sequencing as shown here. ***Previously cytotyped as tetraploid *Camelina* (2n=26) by chromosome squash by Hotton et al., 2020, and found to be tetraploid *Camelina rumelica* (2n=26) by DNA sequencing as shown here. ****Previously cytotyped as hexaploid *Camelina* (2n=38) by chromosome squash by Hotton et al., 2020, and found to be hexaploid *Camelina microcarpa* (2n=38) by DNA sequencing as shown here.

### 3.4 KASP genotyping approach distinguished between functional and subfunctional alleles of *FLC* on chromosome 20

Currently the only genotyping approach to distinguish functional from subfunctional alleles of the *FLC* gene on chromosome 20 is qRT-PCR, which was developed from camelina accessions we shared through an intermediary (Chao et al., 2019). Many plant breeding programs make use of DNA-based Kompetitive Allele Specific PCR (KASP) genotyping, so we aimed to optimize this genotyping approach for this marker. KASP primers were designed for the SNV found in exon 5 of FLC on chromosome 20. While individual primers in this this primer set contained many off target binding sites, these sites were not adjacent to each other and thus would not allow for off target amplification to a degree that could confound results. This KASP primer set was effective in distinguishing winter from spring type camelina. (Figure 3 with specific camelina accessions used in analysis in Supplementary Table 3).

**Figure 3.**
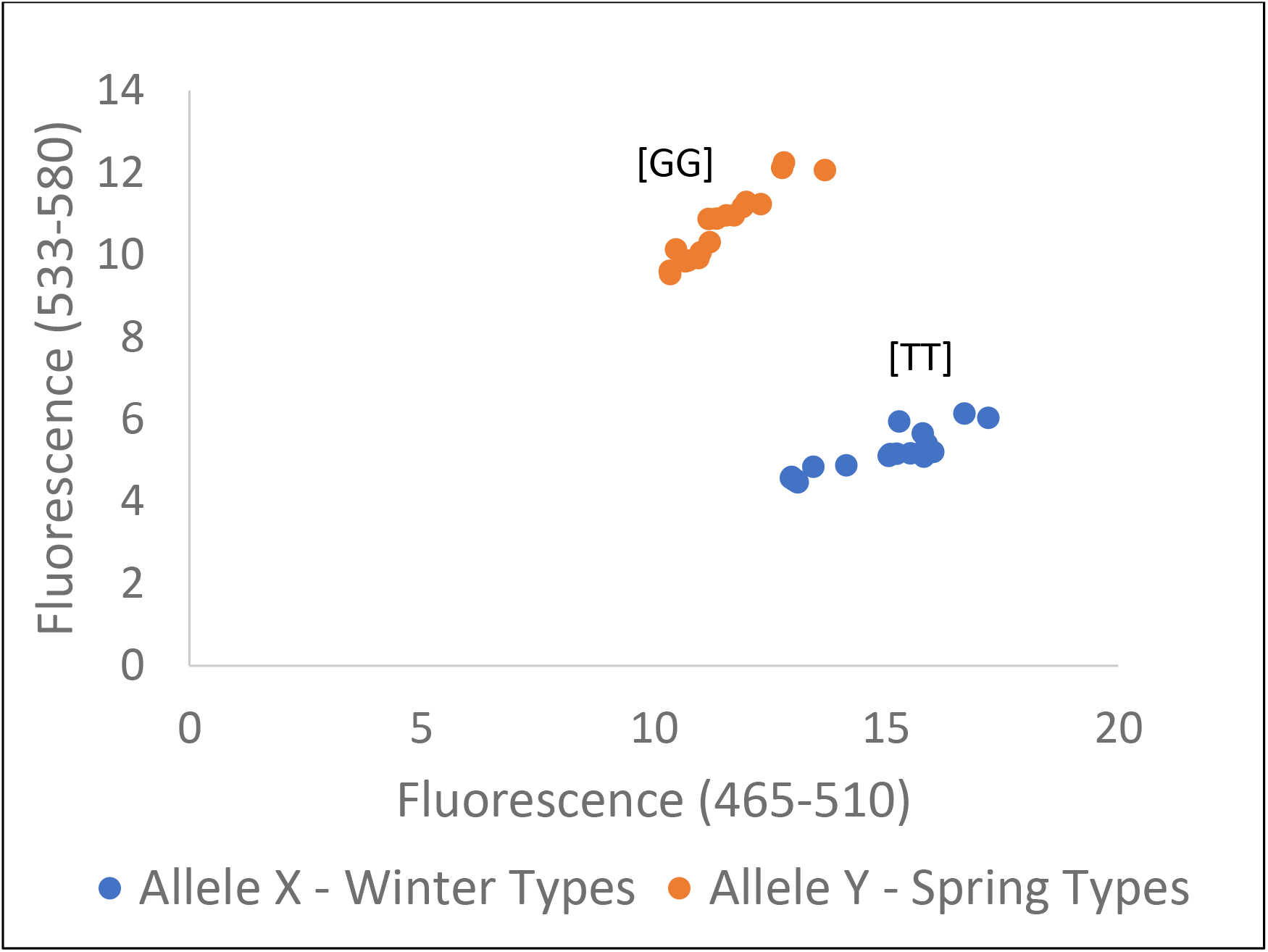
Kompetitive Allele-Specific PCR (KASP) genotyping results that differentiate winter from spring growth habit accessions of *Camelina sativa* using the previously identified single basepair INDEL in exon 5 of the *FLOWERING LOCUS C* gene on chromosome 20 (Anderson et al., 2018). Specific accessions used for this analysis are listed in Supplementary Table 2.

### 3.5 Discovery of facultative winter camelina accession PI650163

The majority of the accessions within our camelina germplasm had a spring-type growth habit (Supplementary Table 1). Based upon our fall 2015 bolting notes, we conducted a follow-up greenhouse study where we planted each accession/line (20) that was scored as slow bolting into pots in triplicate without a vernalizing cold treatment at any point. Fourteen of these accessions did not bolt within the time frame of the experiment. PI 650163 repeated the slow bolting phenotype observed in fall 2015, four bolted at a moderate rate, and one bolted very quickly (Figure 4). The latter may have been subjected to a more stressful micro-environment in fall 2015 resulting in it being mislabeled as slow bolting. By spring of 2016, none of the spring type camelina plants had survived the Minnesota winter.

**Figure 4.**
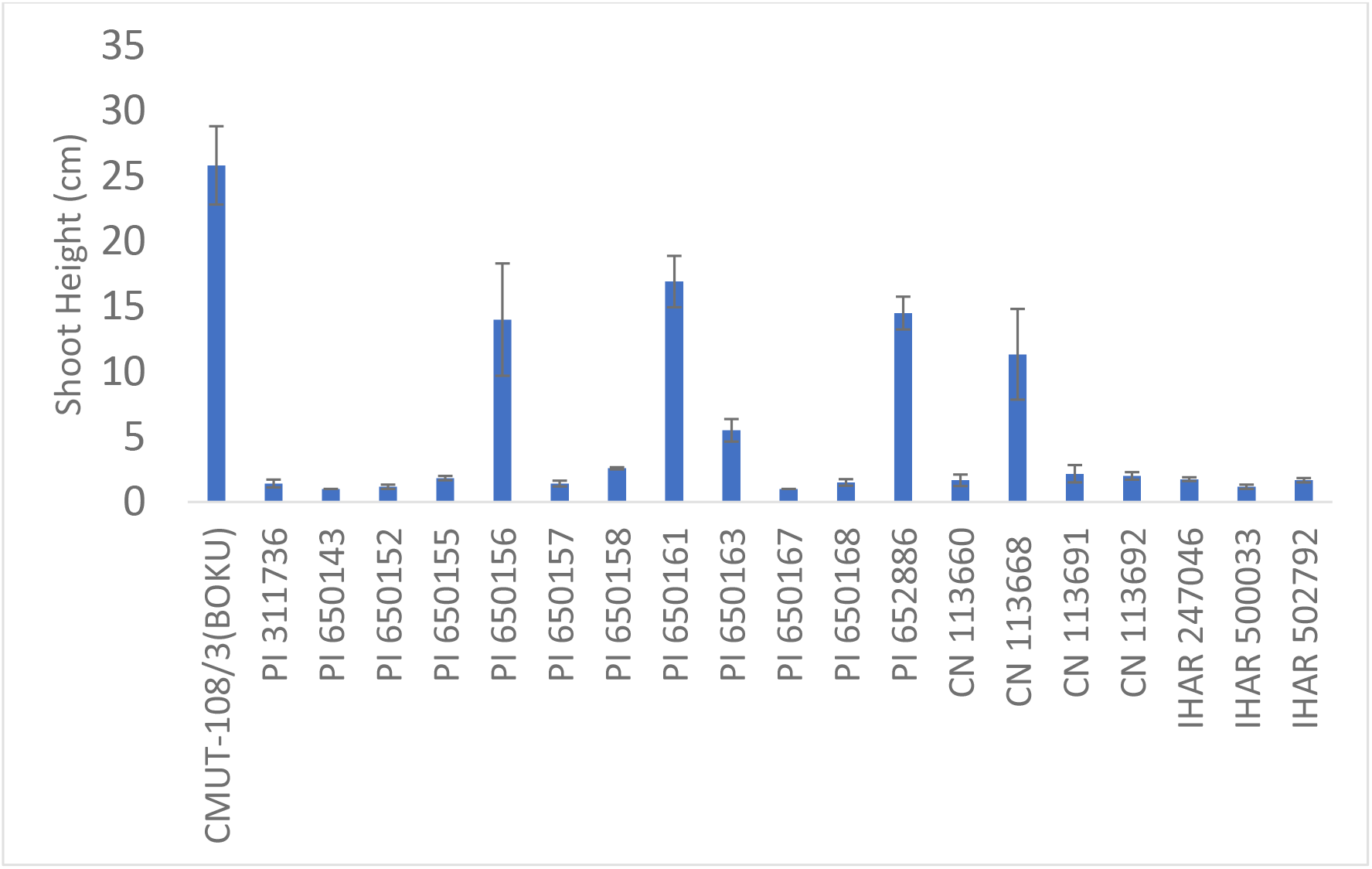
Camelina shoot height observed in a greenhouse 34 days after planting on February, 19^th^ 2016 with no vernalization. Experimental design was a randomized complete block and n=3. The greenhouse temperature was maintained near 20 °C and was supplemented with halogen lighting to ensure 16h light was available each day.

### 3.6 Discovery of winter-hardy, early flowering and maturing camelina accession PI 650163-1

In the fall of 2017, PI650163 was planted in Rosemount, MN with the hypothesis that it may be able to survive a Minnesota winter due to its facultative (slow bolting) winter type growth habit. On May 18^th^, 2018, a single plant from this plot (PI 650163-1) was selected since it was the first plant flowering in this plot as well as in the entire field. PI 650163-1 flowered nine and 17 days before Ames 33292 (‘Joelle’) in 2019 and 2020 in St. Paul, MN, respectively (Table 3). Physiological maturity was also scored in St. Paul, MN in 2020, using the BBCH scale developed by Martinelli and Galasso (2011). PI 650163-1 reached 50% maturity nine days before Ames 33292 (Table 3).

**Table 3.**
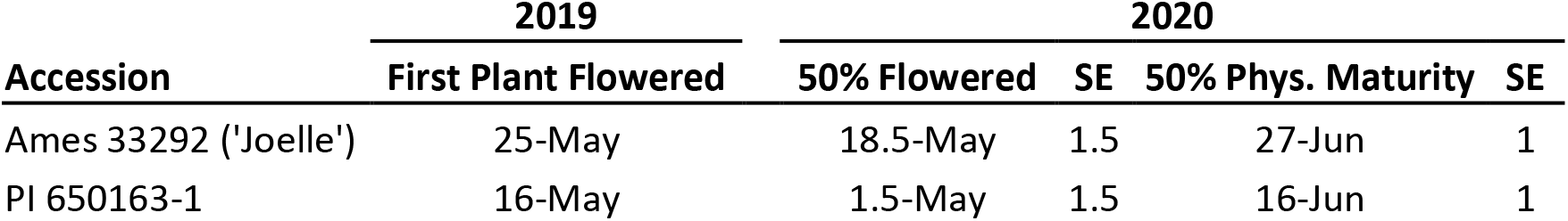
Flowering and maturity visual observations made in the field in St. Paul. Single 1.5 m rows of Camelina were planted on Oct. 18^th^, 2018 (n=1) and Sept. 9^th^, 2019 (n=3), respectively.

| **Accession** | **2019** | **2020** |  |  |  |
| --- | --- | --- | --- | --- | --- |
|  | **First Plant Flowered** | **50% Flowered** | **SE** | **50% Phys. Maturity** | **SE** |
| Ames 33292 ('Joelle') | 25-May | 18.5-May | 1.5 | 27-Jun | 1 |
| PI 650163-1 | 16-May | 1.5-May | 1.5 | 16-Jun | 1 |

In canola, an example of the genetic basis of the facultative winter type is increased expression of a repressor of FLC - the *VENALIZATION INSENSITIVE 3* (*VIN3*) gene, which is typically upregulated after exposure to temperatures near 4°C. In facultative winter type canola, *VIN3* has increased expression even at a temperature of 20°C, far above what typically induces its expression of this gene, and thus baseline *FLC* expression is relatively lower in facultative winter type canola than in winter type (Huang et al., 2021). *Camelina sativa* accession PI 650163 and PI 650163-1 did not have any obvious variants in *VIN3*, but one or more other genes in this pathway may have an allele influencing this phenotype. However, since these facultative winter accessions already have a subfunctional allele of *FLC* on chromosome 20, this floral suppression pathway may be less likely to influence this phenotype. Subfunctional genes in the floral promoter pathway such as *FLOWERING LOCUS T* (*FT*) and *CONSTANS (CO)* could also be influencing this phenotype, but again, these accessions did not have any obvious variants in *FT* or in *CO*.

## 4 Conclusion

While we have gained a much greater understanding of camelina genetics with the extensive whole genome resequencing contribution of Li et al., (2020), our analysis of the sequence data for *FLC* pathway genes has led to additional open questions – primarily, what other gene(s) in the vernalization pathway is/are being suppressed in spring camelina accessions with functional *FLC* chromosome 20 alleles?

The genetic relatedness among camelina accessions showed strong clustering by within the winter and facultative winter types. Possible approaches to leverage the relatedness data could be to use it for selecting spring type parents for introducing more diversity into winter camelina breeding programs as well as to select unique parents to breed for increased seed size. Unique accessions such as PI 650143 can also be further characterized.

The new approach we developed using KASP genotyping to differentiate functional versus subfunctional *FLC* alleles on chromosome 20 is expected to accelerate camelina breeding efforts by making genotyping available on widely used platforms and minimizing costs.

The facultative winter camelina accession we identified – PI 650163-1 – appears to hold promise for cropping systems in the Upper Midwest, since it has demonstrated robust winter hardiness in the site years of this study, but also flowers and matures earlier, allowing the primary summer annual crops to receive full light interception sooner. This accession appears to be at a midpoint between spring and winter camelina accessions on a whole genome level, but it remains unclear if its facultative winter nature is controlled by few or many genes.

## 2.5 Acknowledments

The authors would like to thank Dr. Russ Gesch (USDA-ARS, Morris, MN) for providing important agronomic insights into camelina and for providing seed of accession Ames 33292 (‘Joelle’) before it became available in the NPGS. We would like to thank Dr. Laura Marek – curator of oil seeds crops at North Central Regional Plant Introduction Station (NCRPIS), part of the National Plant Germplasm System (NPGS) – for assistance in establishing a collection of *Camelina sativa* (L. Crantz) accessions at the University of Minnesota. We would like to thank Dr. Johann Vollmann for allowing us to study the camelina lines he developed in his breeding program at the Universität für Bodenkultur Wien (BOKU) in Vienna, Austria. We would also like to thank our undergraduate student employees, Claire Biel and Henry Parks, for their assistance with lab and field work.

## References

Anderson, J. V, Horvath, D.P., Doğramaci, M., Dorn, K.M., Chao, W.S., Watkin, E.E., Hernandez, A.G., Marks, M.D., Gesch, R., 2018. Expression of FLOWERING LOCUS C and a frameshift mutation of this gene on chromosome 20 differentiate a summer and winter annual biotype of Camelina sativa. Plant direct 2, e00060. https://doi.org/10.1002/pld3.60

Brock, J.R., Donmez, A.A., Beistein, M.A., Olsen, K.M., 2018. Phylogenetics of Camelina Crantz. (Brassicaceae) and insights on the origin of gold-of-pleasure (Camelina sativa). Mol. Phylogenet. Evol. 127, 834–842. https://doi.org/10.1016/j.ympev.2018.06.031

Brock, J. R., Mandáková, T., McKain, M., Lysak, M. A., & Olsen, K. M. (2022). Chloroplast phylogenomics in Camelina (Brassicaceae) reveals multiple origins of polyploid species and the maternal lineage of C. sativa. Horticulture Research, 9, uhab050. https://doi.org/10.1093/hr/uhab050

Chao, W.S., Wang, H., Horvath, D.P., Anderson, J. V, 2019. Selection of endogenous reference genes for qRT-PCR analysis in Camelina sativa and identification of FLOWERING LOCUS C allele-specific markers to differentiate summer-and winter-biotypes. Ind. Crops Prod. 129, 495–502. https://doi.org/10.1016/j.indcrop.2018.12.017

Chaudhary, R., Koh, C.S., Kagale, S., Tang, L., Wu, S.W., Lv, Z., Mason, A.S., Sharpe, A.G., Diederichsen, A. and Parkin, I.A., 2020. Assessing diversity in the Camelina genus provides insights into the genome structure of Camelina sativa. G3: Genes, Genomes, Genetics, 10(4), 1297–1308. https://doi.org/10.1534/g3.119.400957

Chaudhary, R., Higgins, E.E., Eynck, C., Sharpe, A. and Parkin, I., 2023. Mapping QTL for vernalization requirement identified adaptive divergence of the candidate gene Flowering Locus C in polyploid Camelina sativa. bioRxiv, 2023-05. https://doi.org/10.1101/2023.05.23.541983

Chen H, Wang T, He X, Cai X, Lin R, Liang J, Wu J, King G, Wang X. 2022. [WWW Document] http://brassicadb.cn/#. Brassicaceae Database (BRAD). Nucleic Acids Res. 50 (*D1*). D1432–D1441. (accessed 3.10.2022).

Crucifer Genome Initiative, n.d. Agriculture and Agri-Food Canada Saskatoon Research and Development Center in collaboration with the Global Institute for Food Security and Dr. Isobel Parkin’s lab. [WWW Document].

Danecek, P., Bonfield, J.K., Liddle, J., Marshall, J., Ohan, V., Pollard, M.O., Whitwham, A., Keane, T., McCarthy, S.A., Davies, R.M., Li, H., 2021. Twelve years of SAMtools and BCFtools. Gigascience 10. https://doi.org/10.1093/gigascience/giab008

Eberle, C.A., Thom, M.D., Nemec, K.T., Forcella, F., Lundgren, J.G., Gesch, R.W., Riedell, W.E., Papiernik, S.K., Wagner, A., Peterson, D.H., Eklund, J.J., 2015. Using pennycress, camelina, and canola cash cover crops to provision pollinators. Ind. Crops Prod. 75, 20–25. https://doi.org/10.1016/j.indcrop.2015.06.026

Gesch, R.W., Cermak, S.C., 2011. Sowing date and tillage effects on fall-seeded camelina in the northern corn belt. Agron. J. 103, 980–987. https://doi.org/10.2134/agronj2010.0485

Hotton, S.K., Kammerzell, M., Chan, R., Hernandez, B.T., Young, H.A., Tobias, C., McKeon, T., Brichta, J., Thomson, N.J., Thomson, J.G., 2020. Phenotypic examination of Camelina sativa (L.) Crantz accessions from the USDA-ARS national genetics resource program. Plants 9, 642. https://doi.org/10.3390/plants9050642

Huang, C.H., Sun, R.R., Hu, Y., Zeng, L.P., Zhang, N., Cai, L.M., Zhang, Q., Koch, M.A., Al-Shehbaz, I., Edger, P.P., Pires, J.C., Tan, D.Y., Zhong, Y., Ma, H., 2016. Resolution of Brassicaceae Phylogeny Using Nuclear Genes Uncovers Nested Radiations and Supports Convergent Morphological Evolution. Mol. Biol. Evol. 33, 394–412. https://doi.org/10.1093/molbev/msv226

Huang, L.Y., Min, Y., Schiessl, S., Xiong, X.H., Jan, H.U., He, X., Qian, W., Guan, C.Y., Snowdon, R.J., Hua, W., Guan, M., Qian, L.W., 2021. Integrative analysis of GWAS and transcriptome to reveal novel loci regulation flowering time in semi-winter rapeseed. PLANT Sci. 310. https://doi.org/10.1016/j.plantsci.2021.110980

Kagale, S., Koh, C.S., Nixon, J., Bollina, V., Clarke, W.E., Tuteja, R., Spillane, C., Robinson, S.J., Links, M.G., Clarke, C., Higgins, E.E., Huebert, T., Sharpe, A.G., Parkin, I.A.P., 2014. The emerging biofuel crop Camelina sativa retains a highly undifferentiated hexaploid genome structure. Nat. Commun. 5. https://doi.org/10.1038/ncomms4706

Li, H., Durbin, R., 2009. Fast and accurate short read alignment with Burrows-Wheeler transform. BIOINFORMATICS 25, 1754–1760. https://doi.org/10.1093/bioinformatics/btp324

Li, H., Hu, X., Lovell, J.T., Grabowski, P.P., Mamidi, S., Chen, C., Amirebrahimi, M., Kahanda, I., Mumey, B., Barry, K., Kudrna, D., Schmutz, J., Lachowiec, J., Lu, C.F., 2021. Genetic dissection of natural variation in oilseed traits of camelina by whole-genome resequencing and QTL mapping. Plant Genome 14. https://doi.org/10.1002/tpg2.20110

Madeira, F., Pearce, M., Tivey, A.R.N., Basutkar, P., Lee, J., Edbali, O., Madhusoodanan, N., Kolesnikov, A., Lopez, R., n.d. Search and sequence analysis tools services from EMBL-EBI in 2022. Nucleic Acids Res. https://doi.org/10.1093/nar/gkac240

Martinelli, T., Galasso, I., 2011. Phenological growth stages of Camelina sativa according to the extended BBCH scale. Ann. Appl. Biol. 158, 87–94. https://doi.org/10.1111/j.1744-7348.2010.00444.x

Ott, M.A., Eberle, C.A., Thom, M.D., Archer, D.W., Forcella, F., Gesch, R.W., Wyse, D.L., 2019. Economics and Agronomics of Relay-Cropping Pennycress and Camelina with Soybean in Minnesota. Agron. J. 111, 1281–1292. https://doi.org/10.2134/agronj2018.04.0277

Samach, A., Onouchi, H., Gold, S.E., Ditta, G.S., Schwarz-Sommer, Z., Yanofsky, M.F., Coupland, G., 2000. Distinct roles of CONSTANS target genes in reproductive development of Arabidopsis. Science (80). 288, 1613–1616. https://doi.org/10.1126/science.288.5471.1613

Sheldon, C.C., Rouse, D.T., Finnegan, E.J., Peacock, W.J., Dennis, E.S., 2000. The molecular basis of vernalization: The central role of FLOWERING LOCUS C (FLC). Proc. Natl. Acad. Sci. U. S. A. 97, 3753–3758. https://doi.org/10.1073/pnas.060023597

Standage, D., 2015. deinterleave fastq file [WWW Document]. Biostars, Bioinforma. Explained.

Weyers, S., Thom, M., Forcella, F., Eberle, C., Matthees, H., Gesch, R., Ott, M., Feyereisen, G., Strock, J., Wyse, D., 2019. Reduced Potential for Nitrogen Loss in Cover Crop-Soybean Relay Systems in a Cold Climate. J. Environ. Qual. 48. https://doi.org/10.2134/jeq2018.09.0350

Wittenberg, A., Anderson, J. V, Berti, M.T., 2020. Crop growth and productivity of winter camelina in response to sowing date in the northwestern Corn Belt of the USA. Ind. Crops Prod. 158, 113036. https://doi.org/10.1016/j.indcrop.2020.113036

Zohary, D., Hopf, M., Weiss, E., 2012. Domestication of Plants in the Old World: The Origin and Spread of Domesticated Plants in Southwest Asia, Europe, and the Mediterranean Basin. OUP Oxford.

