## Supplementary Material for "Facultative winter accession of *Camelina sativa* (L. Crantz) with early maturity contributes to understanding of the role of *FLOWERING LOCUS C* in camelina flowering"

**Table of Contents**

**Supplementary Table 1.** *Camelina sativa* (L. Crantz) accessions received by the University of Minnesota.

**Supplementary Table 2.** Heat map of the mean percentage of bases for a given chromosome with non-zero coverage in 100kb windows.

**Supplementary Table 3.** *Camelina sativa* accessions used for Kompetitive Allele-Specific PCR genotyping to distinguish winter from spring growth habit.

**Supplementary Table 1.** *Camelina sativa* (L. Crantz) accessions received by the University of Minnesota.

| Index Number | Accession ID | Source Repository | Year Received |
| --- | --- | --- | --- |
| 1 | CN 101980 | Plant Gene Resources of Canada | 2015 |
| 2 | CN 101981 | Plant Gene Resources of Canada | 2015 |
| 3 | CN 101982 | Plant Gene Resources of Canada | 2015 |
| 4 | CN 101983 | Plant Gene Resources of Canada | 2015 |
| 5 | CN 101984 | Plant Gene Resources of Canada | 2015 |
| 6 | CN 101985 | Plant Gene Resources of Canada | 2015 |
| 7 | CN 101986 | Plant Gene Resources of Canada | 2015 |
| 8 | CN 101987 | Plant Gene Resources of Canada | 2015 |
| 9 | CN 101988 | Plant Gene Resources of Canada | 2015 |
| 10 | CN 101989 | Plant Gene Resources of Canada | 2015 |
| 11 | CN 101990 | Plant Gene Resources of Canada | 2015 |
| 12 | CN 111330 | Plant Gene Resources of Canada | 2015 |
| 13 | CN 111331 | Plant Gene Resources of Canada | 2015 |
| 14 | CN 111332 | Plant Gene Resources of Canada | 2015 |
| 15 | CN 111333 | Plant Gene Resources of Canada | 2015 |
| 16 | CN 111334 | Plant Gene Resources of Canada | 2015 |
| 17 | CN 111335 | Plant Gene Resources of Canada | 2015 |
| 18 | CN 111336 | Plant Gene Resources of Canada | 2015 |
| 19 | CN 113651 | Plant Gene Resources of Canada | 2015 |
| 20 | CN 113652 | Plant Gene Resources of Canada | 2015 |
| 21 | CN 113653 | Plant Gene Resources of Canada | 2015 |
| 22 | CN 113654 | Plant Gene Resources of Canada | 2015 |
| 23 | CN 113655 | Plant Gene Resources of Canada | 2015 |
| 24 | CN 113656 | Plant Gene Resources of Canada | 2015 |
| 25 | CN 113657* | Plant Gene Resources of Canada | 2015 |
| 26 | CN 113658 | Plant Gene Resources of Canada | 2015 |
| 27 | CN 113659 | Plant Gene Resources of Canada | 2015 |
| 28 | CN 113660* | Plant Gene Resources of Canada | 2015 |
| 29 | CN 113661 | Plant Gene Resources of Canada | 2015 |
| 30 | CN 113662* | Plant Gene Resources of Canada | 2015 |
| 31 | CN 113663 | Plant Gene Resources of Canada | 2015 |
| 32 | CN 113664 | Plant Gene Resources of Canada | 2015 |
| 33 | CN 113665 | Plant Gene Resources of Canada | 2015 |
| 34 | CN 113666 | Plant Gene Resources of Canada | 2015 |
| 35 | CN 113667 | Plant Gene Resources of Canada | 2015 |
| 36 | CN 113668 | Plant Gene Resources of Canada | 2015 |
| 37 | CN 113669 | Plant Gene Resources of Canada | 2015 |
| 38 | CN 113670 | Plant Gene Resources of Canada | 2015 |
| 39 | CN 113671 | Plant Gene Resources of Canada | 2015 |
| 40 | CN 113672 | Plant Gene Resources of Canada | 2015 |
| 41 | CN 113673 | Plant Gene Resources of Canada | 2015 |
| 42 | CN 113674 | Plant Gene Resources of Canada | 2015 |
| 43 | CN 113675 | Plant Gene Resources of Canada | 2015 |
| 44 | CN 113676 | Plant Gene Resources of Canada | 2015 |
| 45 | CN 113677 | Plant Gene Resources of Canada | 2015 |
| 46 | CN 113678 | Plant Gene Resources of Canada | 2015 |
| 47 | CN 113679 | Plant Gene Resources of Canada | 2015 |
| 48 | CN 113680 | Plant Gene Resources of Canada | 2015 |
| 49 | CN 113681 | Plant Gene Resources of Canada | 2015 |
| 50 | CN 113682 | Plant Gene Resources of Canada | 2015 |
| 51 | CN 113683 | Plant Gene Resources of Canada | 2015 |
| 52 | CN 113684 | Plant Gene Resources of Canada | 2015 |
| 53 | CN 113685 | Plant Gene Resources of Canada | 2015 |
| 54 | CN 113686 | Plant Gene Resources of Canada | 2015 |
| 55 | CN 113687 | Plant Gene Resources of Canada | 2015 |
| 56 | CN 113688 | Plant Gene Resources of Canada | 2015 |
| 57 | CN 113689 | Plant Gene Resources of Canada | 2015 |
| 58 | CN 113690 | Plant Gene Resources of Canada | 2015 |
| 59 | CN 113691* | Plant Gene Resources of Canada | 2015 |
| 60 | CN 113692* | Plant Gene Resources of Canada | 2015 |
| 61 | CN 113693 | Plant Gene Resources of Canada | 2015 |
| 62 | CN 113694 | Plant Gene Resources of Canada | 2015 |
| 63 | CN 113695 | Plant Gene Resources of Canada | 2015 |
| 64 | CN 113696 | Plant Gene Resources of Canada | 2015 |
| 65 | CN 113697 | Plant Gene Resources of Canada | 2015 |
| 66 | CN 113698 | Plant Gene Resources of Canada | 2015 |
| 67 | CN 113699 | Plant Gene Resources of Canada | 2015 |
| 68 | CN 113700 | Plant Gene Resources of Canada | 2015 |
| 69 | CN 113701 | Plant Gene Resources of Canada | 2015 |
| 70 | CN 113702 | Plant Gene Resources of Canada | 2015 |
| 71 | CN 113703 | Plant Gene Resources of Canada | 2015 |
| 72 | CN 113704 | Plant Gene Resources of Canada | 2015 |
| 73 | CN 113705 | Plant Gene Resources of Canada | 2015 |
| 74 | CN 113706 | Plant Gene Resources of Canada | 2015 |
| 75 | CN 113707 | Plant Gene Resources of Canada | 2015 |
| 76 | CN 113708 | Plant Gene Resources of Canada | 2015 |
| 77 | CN 113709 | Plant Gene Resources of Canada | 2015 |
| 78 | CN 113710 | Plant Gene Resources of Canada | 2015 |
| 79 | CN 113711 | Plant Gene Resources of Canada | 2015 |
| 80 | CN 113712 | Plant Gene Resources of Canada | 2015 |
| 81 | CN 113713 | Plant Gene Resources of Canada | 2015 |
| 82 | CN 113714 | Plant Gene Resources of Canada | 2015 |
| 83 | CN 113715 | Plant Gene Resources of Canada | 2015 |
| 84 | CN 113716 | Plant Gene Resources of Canada | 2015 |
| 85 | CN 113717 | Plant Gene Resources of Canada | 2015 |
| 86 | CN 113718 | Plant Gene Resources of Canada | 2015 |
| 87 | CN 113719 | Plant Gene Resources of Canada | 2015 |
| 88 | CN 113720 | Plant Gene Resources of Canada | 2015 |
| 89 | CN 113721 | Plant Gene Resources of Canada | 2015 |
| 90 | CN 113722 | Plant Gene Resources of Canada | 2015 |
| 91 | CN 113723 | Plant Gene Resources of Canada | 2015 |
| 92 | CN 113724 | Plant Gene Resources of Canada | 2015 |
| 93 | CN 113725 | Plant Gene Resources of Canada | 2015 |
| 94 | CN 113726 | Plant Gene Resources of Canada | 2015 |
| 95 | CN 113727 | Plant Gene Resources of Canada | 2015 |
| 96 | CN 113728 | Plant Gene Resources of Canada | 2015 |
| 97 | CN 113729 | Plant Gene Resources of Canada | 2015 |
| 98 | CN 113730 | Plant Gene Resources of Canada | 2015 |
| 99 | CN 113731 | Plant Gene Resources of Canada | 2015 |
| 100 | CN 113732 | Plant Gene Resources of Canada | 2015 |
| 101 | CN 113733 | Plant Gene Resources of Canada | 2015 |
| 102 | CN 113734 | Plant Gene Resources of Canada | 2015 |
| 103 | CN 113735 | Plant Gene Resources of Canada | 2015 |
| 104 | CN 113736 | Plant Gene Resources of Canada | 2015 |
| 105 | CN 113737 | Plant Gene Resources of Canada | 2015 |
| 106 | CN 113738 | Plant Gene Resources of Canada | 2015 |
| 107 | CN 113739 | Plant Gene Resources of Canada | 2015 |
| 108 | CN 113740 | Plant Gene Resources of Canada | 2015 |
| 109 | CN 113741 | Plant Gene Resources of Canada | 2015 |
| 110 | CN 113742 | Plant Gene Resources of Canada | 2015 |
| 111 | CN 113743 | Plant Gene Resources of Canada | 2015 |
| 112 | CN 113744 | Plant Gene Resources of Canada | 2015 |
| 113 | CN 113745 | Plant Gene Resources of Canada | 2015 |
| 114 | CN 113746 | Plant Gene Resources of Canada | 2015 |
| 115 | CN 113747 | Plant Gene Resources of Canada | 2015 |
| 116 | CN 113748 | Plant Gene Resources of Canada | 2015 |
| 117 | CN 113749 | Plant Gene Resources of Canada | 2015 |
| 118 | CN 113750 | Plant Gene Resources of Canada | 2015 |
| 119 | CN 113751 | Plant Gene Resources of Canada | 2015 |
| 120 | CN 113752 | Plant Gene Resources of Canada | 2015 |
| 121 | CN 113753 | Plant Gene Resources of Canada | 2015 |
| 122 | CN 113754 | Plant Gene Resources of Canada | 2015 |
| 123 | CN 113755 | Plant Gene Resources of Canada | 2015 |
| 124 | CN 113756 | Plant Gene Resources of Canada | 2015 |
| 125 | CN 113757 | Plant Gene Resources of Canada | 2015 |
| 126 | CN 113758 | Plant Gene Resources of Canada | 2015 |
| 127 | CN 113759 | Plant Gene Resources of Canada | 2015 |
| 128 | CN 113760 | Plant Gene Resources of Canada | 2015 |
| 129 | CN 113761 | Plant Gene Resources of Canada | 2015 |
| 130 | CN 30475 | Plant Gene Resources of Canada | 2015 |
| 131 | CN 30476 | Plant Gene Resources of Canada | 2015 |
| 132 | CN 30477 | Plant Gene Resources of Canada | 2015 |
| 133 | CN 30478 | Plant Gene Resources of Canada | 2015 |
| 134 | CN 30479 | Plant Gene Resources of Canada | 2015 |
| 135 | CN 45816 | Plant Gene Resources of Canada | 2015 |
| 136 | IHAR 247002 | The Plant Breeding & Acclimatization Institute (IHAR), Poland | 2015 |
| 137 | IHAR 247003 | The Plant Breeding & Acclimatization Institute (IHAR), Poland | 2015 |
| 138 | IHAR 247004 | The Plant Breeding & Acclimatization Institute (IHAR), Poland | 2015 |
| 139 | IHAR 247014 | The Plant Breeding & Acclimatization Institute (IHAR), Poland | 2015 |
| 140 | IHAR 247015 | The Plant Breeding & Acclimatization Institute (IHAR), Poland | 2015 |
| 141 | IHAR 247016 | The Plant Breeding & Acclimatization Institute (IHAR), Poland | 2015 |
| 142 | IHAR 247017 | The Plant Breeding & Acclimatization Institute (IHAR), Poland | 2015 |
| 143 | IHAR 247018 | The Plant Breeding & Acclimatization Institute (IHAR), Poland | 2015 |
| 144 | IHAR 247019 | The Plant Breeding & Acclimatization Institute (IHAR), Poland | 2015 |
| 145 | IHAR 247020 | The Plant Breeding & Acclimatization Institute (IHAR), Poland | 2015 |
| 146 | IHAR 247021 | The Plant Breeding & Acclimatization Institute (IHAR), Poland | 2015 |
| 147 | IHAR 247022 | The Plant Breeding & Acclimatization Institute (IHAR), Poland | 2015 |
| 148 | IHAR 247023 | The Plant Breeding & Acclimatization Institute (IHAR), Poland | 2015 |
| 149 | IHAR 247024 | The Plant Breeding & Acclimatization Institute (IHAR), Poland | 2015 |
| 150 | IHAR 247025 | The Plant Breeding & Acclimatization Institute (IHAR), Poland | 2015 |
| 151 | IHAR 247026 | The Plant Breeding & Acclimatization Institute (IHAR), Poland | 2015 |
| 152 | IHAR 247027 | The Plant Breeding & Acclimatization Institute (IHAR), Poland | 2015 |
| 153 | IHAR 247028 | The Plant Breeding & Acclimatization Institute (IHAR), Poland | 2015 |
| 154 | IHAR 247029 | The Plant Breeding & Acclimatization Institute (IHAR), Poland | 2015 |
| 155 | IHAR 247030 | The Plant Breeding & Acclimatization Institute (IHAR), Poland | 2015 |
| 156 | IHAR 247031 | The Plant Breeding & Acclimatization Institute (IHAR), Poland | 2015 |
| 157 | IHAR 247032 | The Plant Breeding & Acclimatization Institute (IHAR), Poland | 2015 |
| 158 | IHAR 247033 | The Plant Breeding & Acclimatization Institute (IHAR), Poland | 2015 |
| 159 | IHAR 247034 | The Plant Breeding & Acclimatization Institute (IHAR), Poland | 2015 |
| 160 | IHAR 247035 | The Plant Breeding & Acclimatization Institute (IHAR), Poland | 2015 |
| 161 | IHAR 247036 | The Plant Breeding & Acclimatization Institute (IHAR), Poland | 2015 |
| 162 | IHAR 247037 | The Plant Breeding & Acclimatization Institute (IHAR), Poland | 2015 |
| 163 | IHAR 247038 | The Plant Breeding & Acclimatization Institute (IHAR), Poland | 2015 |
| 164 | IHAR 247039 | The Plant Breeding & Acclimatization Institute (IHAR), Poland | 2015 |
| 165 | IHAR 247040 | The Plant Breeding & Acclimatization Institute (IHAR), Poland | 2015 |
| 166 | IHAR 247041 | The Plant Breeding & Acclimatization Institute (IHAR), Poland | 2015 |
| 167 | IHAR 247042 | The Plant Breeding & Acclimatization Institute (IHAR), Poland | 2015 |
| 168 | IHAR 247043 | The Plant Breeding & Acclimatization Institute (IHAR), Poland | 2015 |
| 169 | IHAR 247044 | The Plant Breeding & Acclimatization Institute (IHAR), Poland | 2015 |
| 170 | IHAR 247045 | The Plant Breeding & Acclimatization Institute (IHAR), Poland | 2015 |
| 171 | IHAR 247046* | The Plant Breeding & Acclimatization Institute (IHAR), Poland | 2015 |
| 172 | IHAR 247047 | The Plant Breeding & Acclimatization Institute (IHAR), Poland | 2015 |
| 173 | IHAR 247048 | The Plant Breeding & Acclimatization Institute (IHAR), Poland | 2015 |
| 174 | IHAR 247049 | The Plant Breeding & Acclimatization Institute (IHAR), Poland | 2015 |
| 175 | IHAR 247050 | The Plant Breeding & Acclimatization Institute (IHAR), Poland | 2015 |
| 176 | IHAR 247051 | The Plant Breeding & Acclimatization Institute (IHAR), Poland | 2015 |
| 177 | IHAR 247052 | The Plant Breeding & Acclimatization Institute (IHAR), Poland | 2015 |
| 178 | IHAR 247053 | The Plant Breeding & Acclimatization Institute (IHAR), Poland | 2015 |
| 179 | IHAR 247054 | The Plant Breeding & Acclimatization Institute (IHAR), Poland | 2015 |
| 180 | IHAR 247055 | The Plant Breeding & Acclimatization Institute (IHAR), Poland | 2015 |
| 181 | IHAR 247056 | The Plant Breeding & Acclimatization Institute (IHAR), Poland | 2015 |
| 182 | IHAR 247057 | The Plant Breeding & Acclimatization Institute (IHAR), Poland | 2015 |
| 183 | IHAR 247058 | The Plant Breeding & Acclimatization Institute (IHAR), Poland | 2015 |
| 184 | IHAR 500027 | The Plant Breeding & Acclimatization Institute (IHAR), Poland | 2015 |
| 185 | IHAR 500033* | The Plant Breeding & Acclimatization Institute (IHAR), Poland | 2015 |
| 186 | IHAR 500034 | The Plant Breeding & Acclimatization Institute (IHAR), Poland | 2015 |
| 187 | IHAR 500035 | The Plant Breeding & Acclimatization Institute (IHAR), Poland | 2015 |
| 188 | IHAR 500036 | The Plant Breeding & Acclimatization Institute (IHAR), Poland | 2015 |
| 189 | IHAR 500884 | The Plant Breeding & Acclimatization Institute (IHAR), Poland | 2015 |
| 190 | IHAR 502792* | The Plant Breeding & Acclimatization Institute (IHAR), Poland | 2015 |
| 191 | IHAR 502910 | The Plant Breeding & Acclimatization Institute (IHAR), Poland | 2015 |
| 192 | Ames 31231 | US National Plant Germplasm System | 2015 |
| 193 | Ames 33292* | US National Plant Germplasm System | 2015 |
| 194 | PI 258366 | US National Plant Germplasm System | 2015 |
| 195 | PI 258367 | US National Plant Germplasm System | 2015 |
| 196 | PI 304268 | US National Plant Germplasm System | 2015 |
| 197 | PI 304269 | US National Plant Germplasm System | 2015 |
| 198 | PI 304270 | US National Plant Germplasm System | 2015 |
| 199 | PI 304271 | US National Plant Germplasm System | 2015 |
| 200 | PI 311735 | US National Plant Germplasm System | 2015 |
| 201 | PI 311736* | US National Plant Germplasm System | 2015 |
| 202 | PI 597833 | US National Plant Germplasm System | 2015 |
| 203 | PI 633192 | US National Plant Germplasm System | 2015 |
| 204 | PI 633193 | US National Plant Germplasm System | 2015 |
| 205 | PI 633194 | US National Plant Germplasm System | 2015 |
| 206 | PI 650140 | US National Plant Germplasm System | 2015 |
| 207 | PI 650141 | US National Plant Germplasm System | 2015 |
| 208 | PI 650142 | US National Plant Germplasm System | 2015 |
| 209 | PI 650143* | US National Plant Germplasm System | 2015 |
| 210 | PI 650144 | US National Plant Germplasm System | 2015 |
| 211 | PI 650145 | US National Plant Germplasm System | 2015 |
| 212 | PI 650146 | US National Plant Germplasm System | 2015 |
| 213 | PI 650147 | US National Plant Germplasm System | 2015 |
| 214 | PI 650148 | US National Plant Germplasm System | 2015 |
| 215 | PI 650149 | US National Plant Germplasm System | 2015 |
| 216 | PI 650150 | US National Plant Germplasm System | 2015 |
| 217 | PI 650151 | US National Plant Germplasm System | 2015 |
| 218 | PI 650152* | US National Plant Germplasm System | 2015 |
| 219 | PI 650153 | US National Plant Germplasm System | 2015 |
| 220 | PI 650154 | US National Plant Germplasm System | 2015 |
| 221 | PI 650155* | US National Plant Germplasm System | 2015 |
| 222 | PI 650156 | US National Plant Germplasm System | 2015 |
| 223 | PI 650157* | US National Plant Germplasm System | 2015 |
| 224 | PI 650158* | US National Plant Germplasm System | 2015 |
| 225 | PI 650159 | US National Plant Germplasm System | 2015 |
| 226 | PI 650160 | US National Plant Germplasm System | 2015 |
| 227 | PI 650161 | US National Plant Germplasm System | 2015 |
| 228 | PI 650162 | US National Plant Germplasm System | 2015 |
| 229 | PI 650163 | US National Plant Germplasm System | 2015 |
| 230 | PI 650164 | US National Plant Germplasm System | 2015 |
| 231 | PI 650165 | US National Plant Germplasm System | 2015 |
| 232 | PI 650166 | US National Plant Germplasm System | 2015 |
| 233 | PI 650167* | US National Plant Germplasm System | 2015 |
| 234 | PI 650168* | US National Plant Germplasm System | 2015 |
| 235 | PI 652885 | US National Plant Germplasm System | 2015 |
| 236 | PI 652886 | US National Plant Germplasm System | 2015 |
| 237 | CJ7X-1 | Dr. Johann Vollmann, Universität für Bodenkultur Wien (BOKU) | 2015 |
| 238 | CJ9X-1 | Dr. Johann Vollmann, Universität für Bodenkultur Wien (BOKU) | 2015 |
| 239 | CJ10X-1 | Dr. Johann Vollmann, Universität für Bodenkultur Wien (BOKU) | 2015 |
| 240 | CJ11X-1 | Dr. Johann Vollmann, Universität für Bodenkultur Wien (BOKU) | 2015 |
| 241 | CJ12X-1 | Dr. Johann Vollmann, Universität für Bodenkultur Wien (BOKU) | 2015 |
| 242 | CJ13X-1 | Dr. Johann Vollmann, Universität für Bodenkultur Wien (BOKU) | 2015 |
| 243 | CK1X-1 | Dr. Johann Vollmann, Universität für Bodenkultur Wien (BOKU) | 2015 |
| 244 | CJ9X-2 | Dr. Johann Vollmann, Universität für Bodenkultur Wien (BOKU) | 2015 |
| 245 | CJ11X-2 | Dr. Johann Vollmann, Universität für Bodenkultur Wien (BOKU) | 2015 |
| 246 | CK1X-2 | Dr. Johann Vollmann, Universität für Bodenkultur Wien (BOKU) | 2015 |
| 247 | CJ6X-3 | Dr. Johann Vollmann, Universität für Bodenkultur Wien (BOKU) | 2015 |
| 248 | CJ11X-4 | Dr. Johann Vollmann, Universität für Bodenkultur Wien (BOKU) | 2015 |
| 249 | CK1X-4 | Dr. Johann Vollmann, Universität für Bodenkultur Wien (BOKU) | 2015 |
| 250 | CJ9X-7 | Dr. Johann Vollmann, Universität für Bodenkultur Wien (BOKU) | 2015 |
| 251 | CJ12X-7 | Dr. Johann Vollmann, Universität für Bodenkultur Wien (BOKU) | 2015 |
| 252 | CJ13X-7 | Dr. Johann Vollmann, Universität für Bodenkultur Wien (BOKU) | 2015 |
| 253 | CK2X-7 | Dr. Johann Vollmann, Universität für Bodenkultur Wien (BOKU) | 2015 |
| 254 | CK3X-7 | Dr. Johann Vollmann, Universität für Bodenkultur Wien (BOKU) | 2015 |
| 255 | CJ11X-8 | Dr. Johann Vollmann, Universität für Bodenkultur Wien (BOKU) | 2015 |
| 256 | CJ11X-9 | Dr. Johann Vollmann, Universität für Bodenkultur Wien (BOKU) | 2015 |
| 257 | CJ12X-9 | Dr. Johann Vollmann, Universität für Bodenkultur Wien (BOKU) | 2015 |
| 258 | CK2X-9 | Dr. Johann Vollmann, Universität für Bodenkultur Wien (BOKU) | 2015 |
| 259 | CK5X-9 | Dr. Johann Vollmann, Universität für Bodenkultur Wien (BOKU) | 2015 |
| 260 | CJ11X-11 | Dr. Johann Vollmann, Universität für Bodenkultur Wien (BOKU) | 2015 |
| 261 | CJ9X-13 | Dr. Johann Vollmann, Universität für Bodenkultur Wien (BOKU) | 2015 |
| 262 | CJ10X-13 | Dr. Johann Vollmann, Universität für Bodenkultur Wien (BOKU) | 2015 |
| 263 | CJ11X-13 | Dr. Johann Vollmann, Universität für Bodenkultur Wien (BOKU) | 2015 |
| 264 | CK1X-19 | Dr. Johann Vollmann, Universität für Bodenkultur Wien (BOKU) | 2015 |
| 265 | CJ12X-25 | Dr. Johann Vollmann, Universität für Bodenkultur Wien (BOKU) | 2015 |
| 266 | CK1X-25 | Dr. Johann Vollmann, Universität für Bodenkultur Wien (BOKU) | 2015 |
| 267 | CJ6X-28 | Dr. Johann Vollmann, Universität für Bodenkultur Wien (BOKU) | 2015 |
| 268 | CJ11X-31 | Dr. Johann Vollmann, Universität für Bodenkultur Wien (BOKU) | 2015 |
| 269 | CJ12X-38 | Dr. Johann Vollmann, Universität für Bodenkultur Wien (BOKU) | 2015 |
| 270 | CJ6X-42 | Dr. Johann Vollmann, Universität für Bodenkultur Wien (BOKU) | 2015 |
| 271 | CJ11X-42 | Dr. Johann Vollmann, Universität für Bodenkultur Wien (BOKU) | 2015 |
| 272 | CJ11X-43 | Dr. Johann Vollmann, Universität für Bodenkultur Wien (BOKU) | 2015 |
| 273 | CJ11X-52 | Dr. Johann Vollmann, Universität für Bodenkultur Wien (BOKU) | 2015 |
| 274 | CJ11X-55 | Dr. Johann Vollmann, Universität für Bodenkultur Wien (BOKU) | 2015 |
| 275 | CJ11X-59 | Dr. Johann Vollmann, Universität für Bodenkultur Wien (BOKU) | 2015 |
| 276 | CJ11X-65 | Dr. Johann Vollmann, Universität für Bodenkultur Wien (BOKU) | 2015 |
| 277 | CJ11X-69 | Dr. Johann Vollmann, Universität für Bodenkultur Wien (BOKU) | 2015 |
| 278 | CK1X-69 | Dr. Johann Vollmann, Universität für Bodenkultur Wien (BOKU) | 2015 |
| 279 | CK5X-74 | Dr. Johann Vollmann, Universität für Bodenkultur Wien (BOKU) | 2015 |
| 280 | CJ6X-75 | Dr. Johann Vollmann, Universität für Bodenkultur Wien (BOKU) | 2015 |
| 281 | CK5X-75 | Dr. Johann Vollmann, Universität für Bodenkultur Wien (BOKU) | 2015 |
| 282 | CJ6X-78 | Dr. Johann Vollmann, Universität für Bodenkultur Wien (BOKU) | 2015 |
| 283 | CJ10X-79 | Dr. Johann Vollmann, Universität für Bodenkultur Wien (BOKU) | 2015 |
| 284 | CJ11X-79 | Dr. Johann Vollmann, Universität für Bodenkultur Wien (BOKU) | 2015 |
| 285 | CJ12X-79 | Dr. Johann Vollmann, Universität für Bodenkultur Wien (BOKU) | 2015 |
| 286 | CJ12X-80 | Dr. Johann Vollmann, Universität für Bodenkultur Wien (BOKU) | 2015 |
| 287 | CK2X-80 | Dr. Johann Vollmann, Universität für Bodenkultur Wien (BOKU) | 2015 |
| 288 | CJ11X-85 | Dr. Johann Vollmann, Universität für Bodenkultur Wien (BOKU) | 2015 |
| 289 | CJ11X-86 | Dr. Johann Vollmann, Universität für Bodenkultur Wien (BOKU) | 2015 |
| 290 | CJ11X-87 | Dr. Johann Vollmann, Universität für Bodenkultur Wien (BOKU) | 2015 |
| 291 | CK3X-88 | Dr. Johann Vollmann, Universität für Bodenkultur Wien (BOKU) | 2015 |
| 292 | CJ11X-90 | Dr. Johann Vollmann, Universität für Bodenkultur Wien (BOKU) | 2015 |
| 293 | CK1X-92 | Dr. Johann Vollmann, Universität für Bodenkultur Wien (BOKU) | 2015 |
| 294 | CJ11X-92 | Dr. Johann Vollmann, Universität für Bodenkultur Wien (BOKU) | 2015 |
| 295 | CK1X-93 | Dr. Johann Vollmann, Universität für Bodenkultur Wien (BOKU) | 2015 |
| 296 | CK5X-97 | Dr. Johann Vollmann, Universität für Bodenkultur Wien (BOKU) | 2015 |
| 297 | CJ11X-96 | Dr. Johann Vollmann, Universität für Bodenkultur Wien (BOKU) | 2015 |
| 298 | CK1X-98 | Dr. Johann Vollmann, Universität für Bodenkultur Wien (BOKU) | 2015 |
| 299 | CJ11X-97 | Dr. Johann Vollmann, Universität für Bodenkultur Wien (BOKU) | 2015 |
| 300 | CJ11X-98 | Dr. Johann Vollmann, Universität für Bodenkultur Wien (BOKU) | 2015 |
| 301 | CJ11X-100 | Dr. Johann Vollmann, Universität für Bodenkultur Wien (BOKU) | 2015 |
| 302 | CJ11X-101 | Dr. Johann Vollmann, Universität für Bodenkultur Wien (BOKU) | 2015 |
| 303 | CK1X-104 | Dr. Johann Vollmann, Universität für Bodenkultur Wien (BOKU) | 2015 |
| 304 | CJ6X-105 | Dr. Johann Vollmann, Universität für Bodenkultur Wien (BOKU) | 2015 |
| 305 | CK5X-105 | Dr. Johann Vollmann, Universität für Bodenkultur Wien (BOKU) | 2015 |
| 306 | CJ11X-103 | Dr. Johann Vollmann, Universität für Bodenkultur Wien (BOKU) | 2015 |
| 307 | CJ11X-104 | Dr. Johann Vollmann, Universität für Bodenkultur Wien (BOKU) | 2015 |
| 308 | CJ11X-105 | Dr. Johann Vollmann, Universität für Bodenkultur Wien (BOKU) | 2015 |
| 309 | CJ11X-106 | Dr. Johann Vollmann, Universität für Bodenkultur Wien (BOKU) | 2015 |
| 310 | CK1X-110 | Dr. Johann Vollmann, Universität für Bodenkultur Wien (BOKU) | 2015 |
| 311 | CK2X-110 | Dr. Johann Vollmann, Universität für Bodenkultur Wien (BOKU) | 2015 |
| 312 | CK5X-110 | Dr. Johann Vollmann, Universität für Bodenkultur Wien (BOKU) | 2015 |
| 313 | CK5X-111 | Dr. Johann Vollmann, Universität für Bodenkultur Wien (BOKU) | 2015 |
| 314 | CJ11X-109 | Dr. Johann Vollmann, Universität für Bodenkultur Wien (BOKU) | 2015 |
| 315 | CJ9X-115 | Dr. Johann Vollmann, Universität für Bodenkultur Wien (BOKU) | 2015 |
| 316 | CJ11X-110 | Dr. Johann Vollmann, Universität für Bodenkultur Wien (BOKU) | 2015 |
| 317 | CJ13X-115 | Dr. Johann Vollmann, Universität für Bodenkultur Wien (BOKU) | 2015 |
| 318 | CJ11X-111 | Dr. Johann Vollmann, Universität für Bodenkultur Wien (BOKU) | 2015 |
| 319 | CK2X-116 | Dr. Johann Vollmann, Universität für Bodenkultur Wien (BOKU) | 2015 |
| 320 | CK3X-116 | Dr. Johann Vollmann, Universität für Bodenkultur Wien (BOKU) | 2015 |
| 321 | CK5X-116 | Dr. Johann Vollmann, Universität für Bodenkultur Wien (BOKU) | 2015 |
| 322 | CK2X-117 | Dr. Johann Vollmann, Universität für Bodenkultur Wien (BOKU) | 2015 |
| 323 | CK1X-118 | Dr. Johann Vollmann, Universität für Bodenkultur Wien (BOKU) | 2015 |
| 324 | CK2X-118 | Dr. Johann Vollmann, Universität für Bodenkultur Wien (BOKU) | 2015 |
| 325 | CJ11X-114 | Dr. Johann Vollmann, Universität für Bodenkultur Wien (BOKU) | 2015 |
| 326 | CJ7X-121 | Dr. Johann Vollmann, Universität für Bodenkultur Wien (BOKU) | 2015 |
| 327 | CJ11X-115 | Dr. Johann Vollmann, Universität für Bodenkultur Wien (BOKU) | 2015 |
| 328 | CJ12X-121 | Dr. Johann Vollmann, Universität für Bodenkultur Wien (BOKU) | 2015 |
| 329 | CK3X-121 | Dr. Johann Vollmann, Universität für Bodenkultur Wien (BOKU) | 2015 |
| 330 | CJ11X-116 | Dr. Johann Vollmann, Universität für Bodenkultur Wien (BOKU) | 2015 |
| 331 | CK2X-122 | Dr. Johann Vollmann, Universität für Bodenkultur Wien (BOKU) | 2015 |
| 332 | CK3X-122 | Dr. Johann Vollmann, Universität für Bodenkultur Wien (BOKU) | 2015 |
| 333 | CJ6X-123 | Dr. Johann Vollmann, Universität für Bodenkultur Wien (BOKU) | 2015 |
| 334 | CJ11X-117 | Dr. Johann Vollmann, Universität für Bodenkultur Wien (BOKU) | 2015 |
| 335 | CJ13X-123 | Dr. Johann Vollmann, Universität für Bodenkultur Wien (BOKU) | 2015 |
| 336 | CK1X-123 | Dr. Johann Vollmann, Universität für Bodenkultur Wien (BOKU) | 2015 |
| 337 | CK2X-123 | Dr. Johann Vollmann, Universität für Bodenkultur Wien (BOKU) | 2015 |
| 338 | CK4X-123 | Dr. Johann Vollmann, Universität für Bodenkultur Wien (BOKU) | 2015 |
| 339 | CJ11X-118 | Dr. Johann Vollmann, Universität für Bodenkultur Wien (BOKU) | 2015 |
| 340 | CJ13X-124 | Dr. Johann Vollmann, Universität für Bodenkultur Wien (BOKU) | 2015 |
| 341 | CJ6X-125 | Dr. Johann Vollmann, Universität für Bodenkultur Wien (BOKU) | 2015 |
| 342 | CJ11X-119 | Dr. Johann Vollmann, Universität für Bodenkultur Wien (BOKU) | 2015 |
| 343 | CJ12X-126 | Dr. Johann Vollmann, Universität für Bodenkultur Wien (BOKU) | 2015 |
| 344 | CK2X-126 | Dr. Johann Vollmann, Universität für Bodenkultur Wien (BOKU) | 2015 |
| 345 | CJ6X-127 | Dr. Johann Vollmann, Universität für Bodenkultur Wien (BOKU) | 2015 |
| 346 | CJ11X-120 | Dr. Johann Vollmann, Universität für Bodenkultur Wien (BOKU) | 2015 |
| 347 | CK5X-125 | Dr. Johann Vollmann, Universität für Bodenkultur Wien (BOKU) | 2015 |
| 348 | CJ12X-128 | Dr. Johann Vollmann, Universität für Bodenkultur Wien (BOKU) | 2015 |
| 349 | CK4X-127 | Dr. Johann Vollmann, Universität für Bodenkultur Wien (BOKU) | 2015 |
| 350 | CJ11X-122 | Dr. Johann Vollmann, Universität für Bodenkultur Wien (BOKU) | 2015 |
| 351 | CJ12X-129 | Dr. Johann Vollmann, Universität für Bodenkultur Wien (BOKU) | 2015 |
| 352 | CK1X-129 | Dr. Johann Vollmann, Universität für Bodenkultur Wien (BOKU) | 2015 |
| 353 | CK2X-129 | Dr. Johann Vollmann, Universität für Bodenkultur Wien (BOKU) | 2015 |
| 354 | CK3X-129 | Dr. Johann Vollmann, Universität für Bodenkultur Wien (BOKU) | 2015 |
| 355 | CK5X-127 | Dr. Johann Vollmann, Universität für Bodenkultur Wien (BOKU) | 2015 |
| 356 | CJ6X-130 | Dr. Johann Vollmann, Universität für Bodenkultur Wien (BOKU) | 2015 |
| 357 | CJ9X-130 | Dr. Johann Vollmann, Universität für Bodenkultur Wien (BOKU) | 2015 |
| 358 | CJ11X-123 | Dr. Johann Vollmann, Universität für Bodenkultur Wien (BOKU) | 2015 |
| 359 | CJ12X-130 | Dr. Johann Vollmann, Universität für Bodenkultur Wien (BOKU) | 2015 |
| 360 | CJ13X-130 | Dr. Johann Vollmann, Universität für Bodenkultur Wien (BOKU) | 2015 |
| 361 | CJ9X-131 | Dr. Johann Vollmann, Universität für Bodenkultur Wien (BOKU) | 2015 |
| 362 | CK1X-131 | Dr. Johann Vollmann, Universität für Bodenkultur Wien (BOKU) | 2015 |
| 363 | CK4X-130 | Dr. Johann Vollmann, Universität für Bodenkultur Wien (BOKU) | 2015 |
| 364 | CK5X-129 | Dr. Johann Vollmann, Universität für Bodenkultur Wien (BOKU) | 2015 |
| 365 | CK1X-132 | Dr. Johann Vollmann, Universität für Bodenkultur Wien (BOKU) | 2015 |
| 366 | CK2X-132 | Dr. Johann Vollmann, Universität für Bodenkultur Wien (BOKU) | 2015 |
| 367 | Celine | Dr. Johann Vollmann, Universität für Bodenkultur Wien (BOKU) | 2015 |
| 368 | CA13X-17 | Dr. Johann Vollmann, Universität für Bodenkultur Wien (BOKU) | 2015 |
| 369 | CMUT-726/1 | Dr. Johann Vollmann, Universität für Bodenkultur Wien (BOKU) | 2015 |
| 370 | Calena | Dr. Johann Vollmann, Universität für Bodenkultur Wien (BOKU) | 2015 |
| 371 | CMUT-603/4 | Dr. Johann Vollmann, Universität für Bodenkultur Wien (BOKU) | 2015 |
| 372 | CMUT-814/1 | Dr. Johann Vollmann, Universität für Bodenkultur Wien (BOKU) | 2015 |
| 373 | CA2X1S-1 | Dr. Johann Vollmann, Universität für Bodenkultur Wien (BOKU) | 2015 |
| 374 | CA2X2S-26 | Dr. Johann Vollmann, Universität für Bodenkultur Wien (BOKU) | 2015 |
| 375 | CMut67-3 | Dr. Johann Vollmann, Universität für Bodenkultur Wien (BOKU) | 2015 |
| 376 | CMut64-2 | Dr. Johann Vollmann, Universität für Bodenkultur Wien (BOKU) | 2015 |
| 377 | CA9X-21 | Dr. Johann Vollmann, Universität für Bodenkultur Wien (BOKU) | 2015 |
| 378 | CG3X-40 | Dr. Johann Vollmann, Universität für Bodenkultur Wien (BOKU) | 2015 |
| 379 | CA13X_1S-21 | Dr. Johann Vollmann, Universität für Bodenkultur Wien (BOKU) | 2015 |
| 380 | CA4X-4 | Dr. Johann Vollmann, Universität für Bodenkultur Wien (BOKU) | 2015 |
| 381 | CA9X-21 | Dr. Johann Vollmann, Universität für Bodenkultur Wien (BOKU) | 2015 |
| 382 | CA9X-8 | Dr. Johann Vollmann, Universität für Bodenkultur Wien (BOKU) | 2015 |
| 383 | CA13X_2S-24 | Dr. Johann Vollmann, Universität für Bodenkultur Wien (BOKU) | 2015 |
| 384 | CA9X-7 | Dr. Johann Vollmann, Universität für Bodenkultur Wien (BOKU) | 2015 |
| 385 | CA13X-13 | Dr. Johann Vollmann, Universität für Bodenkultur Wien (BOKU) | 2015 |
| 386 | CA13X_1S-23 | Dr. Johann Vollmann, Universität für Bodenkultur Wien (BOKU) | 2015 |
| 387 | M2:4-bulk/1211 | Dr. Johann Vollmann, Universität für Bodenkultur Wien (BOKU) | 2015 |
| 388 | M2:4-bulk/1333 | Dr. Johann Vollmann, Universität für Bodenkultur Wien (BOKU) | 2015 |
| 389 | M2:4-bulk/1006 | Dr. Johann Vollmann, Universität für Bodenkultur Wien (BOKU) | 2015 |
| 390 | CU009 | Dr. Johann Vollmann, Universität für Bodenkultur Wien (BOKU) | 2015 |
| 391 | CW103 | Dr. Johann Vollmann, Universität für Bodenkultur Wien (BOKU) | 2015 |
| 392 | CU011 | Dr. Johann Vollmann, Universität für Bodenkultur Wien (BOKU) | 2015 |
| 393 | CMUT-555/4 | Dr. Johann Vollmann, Universität für Bodenkultur Wien (BOKU) | 2015 |
| 394 | CMUT-408/3 | Dr. Johann Vollmann, Universität für Bodenkultur Wien (BOKU) | 2015 |
| 395 | CMUT-838/2 | Dr. Johann Vollmann, Universität für Bodenkultur Wien (BOKU) | 2015 |
| 396 | CW072 | Dr. Johann Vollmann, Universität für Bodenkultur Wien (BOKU) | 2015 |
| 397 | CMUT-195/4 | Dr. Johann Vollmann, Universität für Bodenkultur Wien (BOKU) | 2015 |
| 398 | CW065 | Dr. Johann Vollmann, Universität für Bodenkultur Wien (BOKU) | 2015 |
| 399 | CMUT-555/1 | Dr. Johann Vollmann, Universität für Bodenkultur Wien (BOKU) | 2015 |
| 400 | CMUT-196/2 | Dr. Johann Vollmann, Universität für Bodenkultur Wien (BOKU) | 2015 |
| 401 | CMUT-814/1 | Dr. Johann Vollmann, Universität für Bodenkultur Wien (BOKU) | 2015 |
| 402 | CA13X_1S-23 | Dr. Johann Vollmann, Universität für Bodenkultur Wien (BOKU) | 2015 |
| 403 | CMUT-195/5 | Dr. Johann Vollmann, Universität für Bodenkultur Wien (BOKU) | 2015 |
| 404 | CW088 | Dr. Johann Vollmann, Universität für Bodenkultur Wien (BOKU) | 2015 |
| 405 | CMUT-119/5 | Dr. Johann Vollmann, Universität für Bodenkultur Wien (BOKU) | 2015 |
| 406 | CW064 | Dr. Johann Vollmann, Universität für Bodenkultur Wien (BOKU) | 2015 |
| 407 | CW068 | Dr. Johann Vollmann, Universität für Bodenkultur Wien (BOKU) | 2015 |
| 408 | CMUT-195/7 | Dr. Johann Vollmann, Universität für Bodenkultur Wien (BOKU) | 2015 |
| 409 | Ersatz: Control/CA13X-2S-44 | Dr. Johann Vollmann, Universität für Bodenkultur Wien (BOKU) | 2015 |
| 410 | CMUT-195/1 | Dr. Johann Vollmann, Universität für Bodenkultur Wien (BOKU) | 2015 |
| 411 | CA13X_1S-15 | Dr. Johann Vollmann, Universität für Bodenkultur Wien (BOKU) | 2015 |
| 412 | C 194 | Dr. Johann Vollmann, Universität für Bodenkultur Wien (BOKU) | 2015 |
| 413 | C 124 | Dr. Johann Vollmann, Universität für Bodenkultur Wien (BOKU) | 2015 |
| 414 | CU007 | Dr. Johann Vollmann, Universität für Bodenkultur Wien (BOKU) | 2015 |
| 415 | CU008 | Dr. Johann Vollmann, Universität für Bodenkultur Wien (BOKU) | 2015 |
| 416 | CMUT-108/3 | Dr. Johann Vollmann, Universität für Bodenkultur Wien (BOKU) | 2015 |
| 417 | CU005 | Dr. Johann Vollmann, Universität für Bodenkultur Wien (BOKU) | 2015 |
| 418 | CW052 | Dr. Johann Vollmann, Universität für Bodenkultur Wien (BOKU) | 2015 |
| 419 | CW091 | Dr. Johann Vollmann, Universität für Bodenkultur Wien (BOKU) | 2015 |
| 420 | CU027 | Dr. Johann Vollmann, Universität für Bodenkultur Wien (BOKU) | 2015 |
| 421 | 403 Arkangelsk | N.I. Vavilov All-Russian Institute of Plant Genetic Resources | 2016 |
| 422 | 407 Karelia | N.I. Vavilov All-Russian Institute of Plant Genetic Resources | 2016 |
| 423 | 404 Arkagelsk | N.I. Vavilov All-Russian Institute of Plant Genetic Resources | 2016 |
| 424 | 234 Kalinin | N.I. Vavilov All-Russian Institute of Plant Genetic Resources | 2016 |
| 425 | 1998 Vologda | N.I. Vavilov All-Russian Institute of Plant Genetic Resources | 2016 |
| 426 | 170 Kalinin | N.I. Vavilov All-Russian Institute of Plant Genetic Resources | 2016 |
| 427 | 257 Tambov | N.I. Vavilov All-Russian Institute of Plant Genetic Resources | 2016 |
| 428 | 4066 Omsk | N.I. Vavilov All-Russian Institute of Plant Genetic Resources | 2016 |
| 429 | 1978 Tatars tan | N.I. Vavilov All-Russian Institute of Plant Genetic Resources | 2016 |
| 430 | 405 Volgograd | N.I. Vavilov All-Russian Institute of Plant Genetic Resources | 2016 |
| 431 | 444 Bashkiria | N.I. Vavilov All-Russian Institute of Plant Genetic Resources | 2016 |
| 432 | 4143 Irkutsk | N.I. Vavilov All-Russian Institute of Plant Genetic Resources | 2016 |
| 433 | 4134 Rostov | N.I. Vavilov All-Russian Institute of Plant Genetic Resources | 2016 |
| 434 | 4052 Omsk | N.I. Vavilov All-Russian Institute of Plant Genetic Resources | 2016 |
| 435 | 78 Kalinin | N.I. Vavilov All-Russian Institute of Plant Genetic Resources | 2016 |
| *Indicates that this accession exhibited delayed flowering and required vernalization to reach maturity – i.e. had the winter growth habit. Discrepancies in growth habit among our observations (Ott et al., 2022), those by Li et al., (2020), and Hotton et al., 2020 are as follows: Ott et al., 2022 and Li et al., (2020) observed that PI 633193 had a spring growth habit, while Hotton et al., (2020) observed it to have a winter growth habit. Ott et al., (2022) and Hotton et al., (2020) observed PI 650167 to have a winter growth habit, while Li et al., 2020 observed it to have a spring growth habit. Ott et al., (2022) and Hotton et al., (2020) observed PI 650157 to have a mix of both spring and winter growth habits, while Li et al., (2020) observed it to have a spring growth habit. Ott et al., (2022) observed CN 113692 (Zarja Socializma Auslese 1 or CS041 in Li et al., (2020) naming convention) to have a winter growth habit, while Li et al., (2020) observed it to have a spring growth habit, and it was not observed by Hotton et al., (2020). | | | |

**Supplementary Table 2.** Heat map of the mean percentage of bases for a given chromosome with non-zero coverage in 100kb windows from whole genome sequenced paired end reads aligned to the *Camelina sativa* reference genome. *Camelina spp*. accessions PI650135* (diploid *Camelina neglecta*), PI650133* (diploid *Camelina hispida*), and USDA-NPGS stated *Camelina sativa* accessions PI258367**, PI650141**, PI650143**, PI650144**, PI650146**, PI650164**, PI650152*** (*Camelina rumelica* 2n=26), CN113657 (*Camelina rumelica* 2n=26), PI650167**** (*Camelina microcarpa* 2n=38), IHAR50003 (*Camelina microcarpa* 2n=38), PI650168**, PI650163-1. *Previously cytotyped as diploid *C. spp.* by DNA sequencing by Chaudhary et al., 2020. **Previously cytotyped as hexaploid *Camelina* (2n=38) by chromosome squash by Hotton et al., 2020, but found to be hexaploid *Camelina sativa* (2n=40) by DNA sequencing as shown here. ***Previously cytotyped as tetraploid *Camelina* (2n=26) by chromosome squash by Hotton et al., 2020, and found to be tetraploid *Camelina rumelica* (2n=26) by DNA sequencing as shown here. ****Previously cytotyped as hexaploid *Camelina* (2n=38) by chromosome squash by Hotton et al., 2020, and found to be hexaploid *Camelina microcarpa* (2n=38) by DNA sequencing as shown here.

| **Chromosome** | **Subgenome** | **PI650135** | **PI650133** | **PI258367** | **PI650141** | **PI650143** | **PI650144** | **PI650146** | **PI650164** | **PI650152** | **CN113657** | **PI650167** | **IHAR50003** | **PI650168** | **PI650163-1** |
| --- | --- | --- | --- | --- | --- | --- | --- | --- | --- | --- | --- | --- | --- | --- | --- |
| Chr4 | 1 | 0.825 | 0.667 | 0.932 | 0.929 | 0.933 | 0.931 | 0.932 | 0.931 | 0.815 | 0.807 | 0.933 | 0.930 | 0.894 | 0.934 |
| Chr7 | 1 | 0.844 | 0.653 | 0.928 | 0.924 | 0.929 | 0.927 | 0.922 | 0.929 | 0.821 | 0.813 | 0.932 | 0.928 | 0.887 | 0.929 |
| Chr8 | 1 | 0.853 | 0.666 | 0.936 | 0.930 | 0.930 | 0.932 | 0.930 | 0.933 | 0.823 | 0.815 | 0.933 | 0.931 | 0.894 | 0.930 |
| Chr11 | 1 | 0.854 | 0.669 | 0.938 | 0.932 | 0.933 | 0.934 | 0.933 | 0.932 | 0.822 | 0.814 | 0.933 | 0.929 | 0.897 | 0.933 |
| Chr14 | 1 | 0.840 | 0.678 | 0.929 | 0.918 | 0.924 | 0.921 | 0.922 | 0.921 | 0.822 | 0.814 | 0.923 | 0.918 | 0.869 | 0.924 |
| Chr19 | 1 | 0.852 | 0.652 | 0.924 | 0.922 | 0.925 | 0.924 | 0.923 | 0.925 | 0.813 | 0.805 | 0.925 | 0.923 | 0.886 | 0.924 |
| Chr1 | 2 | 0.579 | 0.622 | 0.933 | 0.932 | 0.931 | 0.934 | 0.932 | 0.934 | 0.744 | 0.727 | 0.932 | 0.927 | 0.900 | 0.930 |
| Chr3 | 2 | 0.568 | 0.644 | 0.932 | 0.927 | 0.928 | 0.929 | 0.930 | 0.930 | 0.746 | 0.729 | 0.929 | 0.927 | 0.892 | 0.928 |
| Chr6 | 2 | 0.571 | 0.637 | 0.938 | 0.936 | 0.941 | 0.938 | 0.938 | 0.938 | 0.750 | 0.734 | 0.936 | 0.931 | 0.904 | 0.940 |
| Chr10 | 2 | 0.574 | 0.637 | 0.938 | 0.938 | 0.939 | 0.939 | 0.936 | 0.935 | 0.744 | 0.727 | 0.936 | 0.932 | 0.900 | 0.939 |
| Chr13 | 2 | 0.578 | 0.640 | 0.936 | 0.935 | 0.934 | 0.935 | 0.936 | 0.936 | 0.752 | 0.736 | 0.935 | 0.929 | 0.903 | 0.933 |
| Chr16 | 2 | 0.583 | 0.631 | 0.940 | 0.927 | 0.937 | 0.930 | 0.929 | 0.931 | 0.755 | 0.740 | 0.935 | 0.931 | 0.897 | 0.937 |
| Chr18 | 2 | 0.576 | 0.629 | 0.938 | 0.936 | 0.938 | 0.938 | 0.936 | 0.937 | 0.745 | 0.728 | 0.937 | 0.934 | 0.901 | 0.938 |
| Chr2 | 3 | 0.492 | 0.826 | 0.926 | 0.926 | 0.927 | 0.928 | 0.926 | 0.926 | 0.814 | 0.806 | 0.742 | 0.648 | 0.891 | 0.927 |
| Chr5 | 3 | 0.490 | 0.818 | 0.924 | 0.913 | 0.919 | 0.919 | 0.915 | 0.916 | 0.792 | 0.781 | 0.733 | 0.643 | 0.873 | 0.918 |
| Chr9 | 3 | 0.482 | 0.824 | 0.922 | 0.915 | 0.914 | 0.917 | 0.919 | 0.915 | 0.801 | 0.791 | 0.739 | 0.648 | 0.882 | 0.914 |
| Chr12 | 3 | 0.487 | 0.820 | 0.922 | 0.908 | 0.913 | 0.910 | 0.910 | 0.916 | 0.815 | 0.806 | 0.738 | 0.649 | 0.875 | 0.912 |
| Chr15 | 3 | 0.490 | 0.827 | 0.925 | 0.917 | 0.920 | 0.920 | 0.919 | 0.922 | 0.819 | 0.809 | 0.747 | 0.657 | 0.882 | 0.919 |
| Chr17 | 3 | 0.495 | 0.813 | 0.926 | 0.917 | 0.920 | 0.920 | 0.918 | 0.921 | 0.807 | 0.797 | 0.763 | 0.677 | 0.885 | 0.919 |
| Chr20 | 3 | 0.476 | 0.819 | 0.923 | 0.918 | 0.916 | 0.920 | 0.919 | 0.919 | 0.804 | 0.793 | 0.733 | 0.641 | 0.884 | 0.916 |

**Supplementary Table 3.** *Camelina sativa* accessions used for Kompetitive Allele-Specific PCR genotyping to distinguish winter from spring growth habit.

| Accession-Rep | Growth Habit | Seed Source | Fluorescence  (465-510) | Fluorescence  (533-580) |
| --- | --- | --- | --- | --- |
| Ames 33292-1 | Winter | Dr. Russ Gesch, USDA-ARS, Morris, MN | 16.023 | 5.203 |
| Ames 33292-2 | Winter | Dr. Russ Gesch, USDA-ARS, Morris, MN | 15.82 | 5.087 |
| Ames 33292-3 | Winter | Dr. Russ Gesch, USDA-ARS, Morris, MN | 13.101 | 4.464 |
| CN 113691-1 | Winter | Plant Gene Resources of Canada | 13.081 | 4.507 |
| CN 113691-2 | Winter | Plant Gene Resources of Canada | 15.093 | 5.153 |
| CN 113691-3 | Winter | Plant Gene Resources of Canada | 15.791 | 5.657 |
| IHAR 502792-1 | Winter | The Plant Breeding & Acclimatization Institute (IHAR), Poland | 12.974 | 4.595 |
| IHAR 502792-2 | Winter | The Plant Breeding & Acclimatization Institute (IHAR), Poland | 12.952 | 4.567 |
| IHAR 502792-3 | Winter | The Plant Breeding & Acclimatization Institute (IHAR), Poland | 13.029 | 4.515 |
| PI 650155-1 | Winter | US National Plant Germplasm System | 15.233 | 5.161 |
| PI 650155-2 | Winter | US National Plant Germplasm System | 15.06 | 5.104 |
| PI 650155-3 | Winter | US National Plant Germplasm System | 15.526 | 5.165 |
| PI 650157-1 | Winter | US National Plant Germplasm System | 17.205 | 6.034 |
| PI 650157-2 | Winter | US National Plant Germplasm System | 14.145 | 4.881 |
| PI 650157-3 | Winter | US National Plant Germplasm System | 13.438 | 4.846 |
| PI 650158-1 | Winter | US National Plant Germplasm System | 15.289 | 5.949 |
| PI 650158-2 | Winter | US National Plant Germplasm System | 16.688 | 6.143 |
| PI 650158-3 | Winter | US National Plant Germplasm System | 15.877 | 5.4 |
| CJ12X-129-1 | Spring | Dr. Johann Vollmann, Universität für Bodenkultur Wien (BOKU) | 11.21 | 10.313 |
| CK5X-9-1 | Spring | Dr. Johann Vollmann, Universität für Bodenkultur Wien (BOKU) | 10.962 | 9.925 |
| CK5X-9-2 | Spring | Dr. Johann Vollmann, Universität für Bodenkultur Wien (BOKU) | 12.766 | 12.119 |
| CK5X-75-1 | Spring | Dr. Johann Vollmann, Universität für Bodenkultur Wien (BOKU) | 13.69 | 12.063 |
| CK5X-97-1 | Spring | Dr. Johann Vollmann, Universität für Bodenkultur Wien (BOKU) | 11.188 | 10.87 |
| CK5X-97-2 | Spring | Dr. Johann Vollmann, Universität für Bodenkultur Wien (BOKU) | 12.808 | 12.246 |
| CN 113704-1 | Spring | Plant Gene Resources of Canada | 10.679 | 9.841 |
| CN 113704-2 | Spring | Plant Gene Resources of Canada | 10.736 | 9.862 |
| CN 113704-3 | Spring | Plant Gene Resources of Canada | 11.014 | 10.062 |
| CN 113746-1 | Spring | Plant Gene Resources of Canada | 11.993 | 11.286 |
| CN 113746-2 | Spring | Plant Gene Resources of Canada | 12.31 | 11.231 |
| CN 113746-3 | Spring | Plant Gene Resources of Canada | 11.356 | 10.883 |
| CN 113746-4 | Spring | Plant Gene Resources of Canada | 11.916 | 11.164 |
| CU008-1 | Spring | Dr. Johann Vollmann, Universität für Bodenkultur Wien (BOKU) | 11.563 | 10.96 |
| CU008-2 | Spring | Dr. Johann Vollmann, Universität für Bodenkultur Wien (BOKU) | 10.483 | 10.132 |
| Vavilov 1998 Vologda-1 | Spring | N.I. Vavilov All-Russian Institute of Plant Genetic Resources | 11.733 | 10.958 |
| Vavilov 1998 Vologda-2 | Spring | N.I. Vavilov All-Russian Institute of Plant Genetic Resources | 10.358 | 9.536 |
| Vavilov 1998 Vologda-3 | Spring | N.I. Vavilov All-Russian Institute of Plant Genetic Resources | 10.351 | 9.613 |
